# Analyzing Biomarker Discovery: Estimating the Reproducibility of Biomarker Sets

**DOI:** 10.1101/2021.05.21.445109

**Authors:** Amir Forouzandeh, Alex Rutar, Sunil V Kalmady, Russell Greiner

## Abstract

Many researchers try to understand a biological condition by identifying *biomarkers*. This is typically done using univariate hypothesis testing over a labeled dataset, declaring a feature to be a biomarker if there is a significant statistical difference between its values for the subjects with different outcomes. However, such sets of proposed biomarkers are often not reproducible – subsequent studies often fail to identify the same sets. Indeed, there is often a very small overlap between the biomarkers proposed in pairs of related studies that explore the same phenotypes over the same distribution of subjects. This paper first defines the *Reproducibility Score* for a labeled dataset as a measure (taking values between 0 and 1) of the reproducibility of the results produced by an arbitrary fixed biomarker discovery process for a given distribution of subjects. We then provide ways to reliably estimate this score by defining algorithms that produce an over-bound and an under-bound for this score for a given dataset and biomarker discovery process, for the case of univariate hypothesis testing on dichotomous groups. We confirm that these approximations are meaningful by providing empirical results on a large number of datasets and show that these predictions match known reproducibility results. We have also created a publicly available website, hosted at https://biomarker.shinyapps.io/BiomarkerReprod/, that produces these *Reproducibility Score* approximations for any given dataset (with continuous or discrete features and binary class labels).

## 1 Introduction

Improved understanding of a disease can lead to better diagnosis and treatment. This often begins by finding “biomarkers”, which is a generic term referring to “a characteristic that is objectively measured and evaluated as an indicator of normal biological processes, pathogenic processes, or pharmacologic responses to a therapeutic intervention”^1^. Typically, these are individual features (*e.g*., expression values of specific genes^2,3^) that follow different distributions (*e.g*., have different mean values) in diseased patients versus healthy controls.

Sometimes, biomedical researchers can identify candidates for biomarkers based on their existing knowledge of the disease etiology and/or cellular pathways. This is done by seeking features that are causally related to the disease (*e.g*., phenylketonuria is caused by mutations in a single gene PAH^4^) or a symptom of it (*e.g*., Hemoglobin A1C for monitoring the degree of glucose metabolism in diabetes^5^). This paper, however, focuses on the usage of statistical tools to discover and evaluate biomarkers. A typical dataset can be viewed as a matrix whose rows each correspond to a subject (say a person) and each column corresponds to a feature (*e.g*., clinical measure, or the expression value of a gene), and with a final column providing the outcome (*e.g*., a binary outcome distinguishing case versus control); see Figure 1.

**Figure 1.**
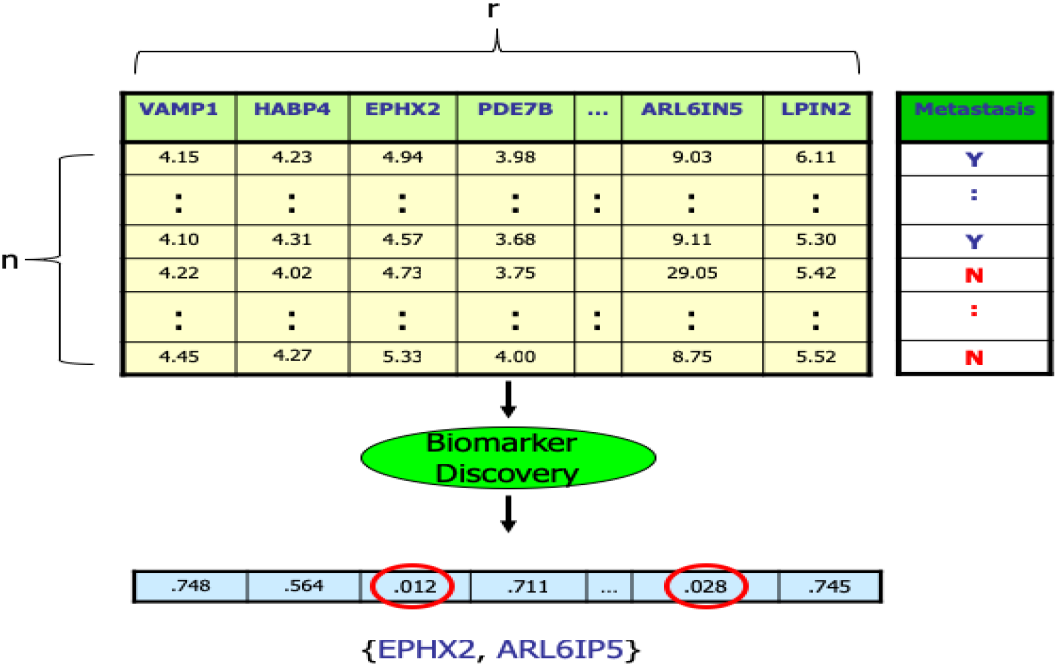
Data matrix, showing t-test p-values for each (shown) feature for the GSE 7390 dataset^14^, with respect to the group outcome (here “Metastasis” for breast cancer); the circled features, with *p <*0.05, are (purported) biomarkers. For notation: We will refer to each of the first *r* columns of the matrix as a “feature”; these are often called “(independent) variables”. We refer to the final column as a “outcome” – *e.g*., case versus control – these are often called “labels”, “dependent variables”, “groups”, “phenotypes” or “classes”. Finally, we will use “subject” to refer to each row of that matrix; these are sometimes called “instances” or “samples”.

These “biomarker discovery studies” (also known as “association studies”) then attempt to determine which of the features (columns) differ significantly among distinct outcomes. Two standard examples of such studies are the “Genome Wide Association Study” (GWAS), over a set of SNPs^6^; and the “Gene Signature Study”, over gene expression values^7^. Typically, this involves first computing a representative statistic for each feature (*e.g*., for continuous entries, running a t-test based on the mean and variance of the case versus control), and then declaring a feature to be a biomarker if the resulting MCC-corrected *p*-value is below 0.05^8^. Here, we use MCC (Multiple Comparison Corrections) as a generic term, which includes both False Discovery Rate (FDR) correction, and Family-Wise Error (FWE) Correction. (Appendix A.2 discusses some of the subtleties here, especially with respect to features.) In some situations, the researchers then validate these biomarkers using a biological or medical process (*e.g*., based on knock-out or amplification studies^9,10^). Other studies validate the proposed biomarkers based on existing biological knowledge. A third class of projects instead use the potential biomarkers to create a computational model – perhaps to learn a classifier^11–13^ – and then measure that down-stream model (perhaps based on its accuracy on a held-out set) and declare the biomarkers to be useful if that model scores well.

A great many papers, however, simply publish the list of purported biomarkers without providing validation for this set; see Appendix **??**. This paper focuses specifically on this case. We address this limitation by providing a falsifiable (statistical) claim about such sets of biomarkers, which suggests a way to validate the proposed biomarker sets.

While some biomarkers are causally related to the associated outcome, this can be difficult to establish (often requiring instrumented studies^15^); but fortunately, in many situations, it may be sufficient for the features to be *correlated* with the phenotype. Here, an ideal biomarker discovery process would identify all-and-only the features that are *consistently* correlated with the associated disease, in that its presence (or absence or …) alone supports that disease. This argues that a proposed biomarker is good if it was *reproducible* – *i.e*., that the biomarkers found in one study, would appear in many (ideally, all) future studies that explore this disease.

This has motivated the use of independent test sets to check the validity of the earlier findings. Unfortunately, many papers report this is not the case – *i.e*., that relatively few biomarkers appear across multiple studies. For example, while the breast cancer studies by van’t Veer *et al*. ^2^ (resp., Wang *et al*. ^3^) reported signatures with 70 (resp., 76) genes, these two sets had only 3 genes in common. Ein-Dor *et al*. ^16^ notes this in another situation: “Only 17 genes appeared in both the list of 456 genes of Sorlie *et al*. ^17^ and the 231 genes of van’t Veer *et al*. ^2^; merely 2 genes were shared between the sets of Sorlie *et al*. and Ramaswamy *et al*. ^18^. Such disparity is not limited to breast cancer but characterizes other human disease datasets (Lossos *et al*. ^19^) such as schizophrenia (Miklos and Maleszka^20^)”. Many others^21–23^ report similar findings. Indeed, Begley and Ellis^24^ report that only 6 of 53 published findings in cancer biology could be confirmed; which Wen *et al*. ^25^ notes is “a rate approaching an alarmingly low 10% of reproducibility”. Moreover, a 2016 Nature survey^26^, of over 1500 scientists, found that 70% of researchers have tried but failed to reproduce another scientist’s experiments, and 52% thought there was a significant ‘crisis’ of reproducibility.

There are many possible reasons for this.

1. Each study should consider the *same well-defined “distribution” over instances* – *e.g*., over the same distribution of ages and genders, etc. If the study attempts to distinguish case from control, then the two sub-populations should only differ in a single characteristic. Unfortunately, matching cases and controls over all possible features is often not achievable.
2. A second issue is with the precise notion of what “*reproducible*” means. Is it a property of a *specific biomarker*, or of *a set of biomarkers*? There is no clear choice for an optimal objective measure. This is especially problematic when dealing with multi-factorial diseases, where the outcomes corresponds to a disjunction over many sub-diseases^27^.
3. A final important issue is the impact of *sample size*. Many studies have a relatively low number of subjects, which increases the probability of finding both false negatives and false positives.

Our analysis assumes that the researchers have addressed (1) by running carefully designed, well-specified studies. Further, we also assume that there is no uncertainty in the outcomes with respect to its clinical or biological definition. We will provide a precise measure of reproducibility (2), as well as some specific implementations, and show empirically how this measurement varies with sample size (3).

In this paper, we assume that biomarkers are stand-alone features. Note each feature could be a pre-defined combination of single features (*e.g*., the average expression values of the genes associated with a pre-defined signalling pathway – see gene enrichment^28^) or networks of genes associated with high loadings of principal component and univariate Pearson correlation values (see PC-corr^29^); but we are not considering *learning* combinations. By a *Biomarker Discovery* process BD(·), we mean a function that takes as input a data matrix of *n* subjects over a set of *r* features and identifies a subset of proposed biomarkers; see Figure 1. We will formally define a *Reproducibility Score*, RS(*D*, BD), to quantify the “reproducibility” of the set of proposed biomarkers produced by the biomarker discovery process: *viz*.,

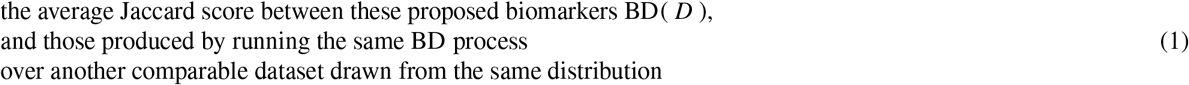

where two datasets are comparable if they have the same number of subjects from each outcome. In order to estimate the reproducibility score in practice, we construct two approximations: an overbound and an underbound. We then provide empirical tests over many datasets, with a focus on *t*-tests as the main biomarker discovery process. We provide many examples for both microarray data (with continuous values) and SNP data (with discrete values) to provide practical evidence for the effectiveness of these approximations. Researchers can use this framework to estimate the reproducibility of the potential results of their biomarker discover study. A low reproducibility score suggests that these biomarkers may not be accurate, potentially because the dataset used is too small, the dataset is too heterogenous, or the biomarker discovery algorithm is not suitable for the dataset. We have also produced a publicly available website, hosted at https://biomarker.shinyapps.io/BiomarkerReprod/, that, given a labeled dataset, computes these estimates of the Reproducibility Score, with respect to any of a variety of biomarker discovery algorithms.

**Outline:** Section 2 formally defines the Reproducibility Score (RS) and describes the challenges of estimating this measure. We then define two approximations for RS: an overbound and an underbound. It also describes some of the standard biomarker discovery algorithms. Section 3 describes extensive empirical studies over many datasets – microarray and mRNAseq data (with continuous values) and SNP data (with discrete values), focusing on a standard Biomarker Discovery process BD(·), to confirm the effectiveness of these approximations. Section 4 summarizes some future work and the contributions of this paper. In the Supplementary Information, Appendix A discusses various notes: a glossary of the various technical terms used, how Biomarker Discovery differs from standard (supervised) Machine Learning, how a combination of a pair of features might be important for a predictive task, even if neither, by itself, is important (towards explaining why it can be so difficult to find biomarkers); etc. Appendix B presents results from other empirical studies, which explore how the RS varies with the type of MCC correction used (including “none”), the p-value threshold, the size of the dataset and the number of iterations of the approximation algorithms. Finally, the approximations we present are motivated by two heuristics (Heuristics 7 and 9). Appendix C presents arguments that motivate these heuristics, and also provide additional empirical evidence that support them.

We close this section by motivating the need for an objective measure for evaluating the quality of a set of biomarkers (Subsection 1.1), then overviewing some earlier studies that discuss the issue of reproducibility in biomarker discovery and/or provide approaches that could be beneficial when dealing with such problems (Subsection 1.2).

### 1.1 Motivation for Evaluating Biomarker Sets

To motivate the need for evaluating association studies, consider first *predictive studies*, which use a labeled dataset, like the one shown at the top of Figure 1, to produce a predictive model (perhaps a decision tree, or a linear classifier) that can be used to classify future subjects – here, into one of the two classes: Y or N. In addition to the learned classifier, the researchers will also compute *a meaningful estimate of its quality* – *i.e*., of the accuracy (or AUROC, Kappa Score, etc.) of this classifier on an independent hold-out set^30^, or the results of *k*-fold cross-validation over the training sample.

By contrast, many association studies report only a set of purported biomarkers, but provide no falsifiable claim about the accuracy of these biomarkers. Many meta-reviews claim that a set of biomarkers is problematic if they are not reproduced in subsequent studies^16,31–33^. Given that biomarkers should be reproducible, we propose evaluating a biomarker set based on its reproducibility score. An accurate estimate of this score can help in at least the following three ways:

1. Researchers can compare various different “comparable” biomarker discovery algorithms to see which produces the biomarker set that is most reproducible. Here, “comparable” corresponds to the standard practice of only considering discovery tools that impose the same criterion, such as the same *p*-value, or only considering features that exhibit the same minimum fold-change. This type of analysis may help to determine errors within the biomarker discovery process. Moreover, we will see that MCC-correction, while useful in removing false-positives, can be detrimental to the goal of producing reproducible biomarkers; similarly, there is no reason to insist on *p <* 0.05 for the statistic test used.
2. A low reproducibility score suggests that few of the proposed biomarkers will be found in another dataset, and highlights the potential that these proposed biomarkers may not be accurate. This could motivate researchers to consider a dataset that is larger, to focus on a more homogenous population, or perhaps consider another biomarker discovery technique.
3. Finally, there are many meta-reviews^21,34,35^ that note the lack of repeatability in many biomarker discovery papers, and question whether the techniques used are to blame. One way to address this concern is to require that each published paper include both the purported set of biomarkers, and also an estimate of its reproducibility score. The same way a prediction study’s “5-fold cross validation” accuracy tells the reader how accurate the classification model should be on new data, this biomarker-discovery reproducibility score will inform the reader whether to expect another study, on a similar dataset, will find many of the same biomarkers. Note that we should view the reproducibility score as necessary for considering a proposed model, but not sufficient – *i.e*., it might rule-out a proposed discovery model, but should not be enough to rule-in a model.

For these reasons, we provide an easy-to-use, publicly available webapp https://biomarker.shinyapps.io/BiomarkerReprod/ that anyone can use to produce meaningful estimates of the reproducibility of a set of biomarkers.

### 1.2 Related Work

There have been many pairs of studies that have each produced biomarkers for the same disease or condition, but found little or no overlap between the two lists of purported biomarkers. Many papers have discussed this issue – some describing this problem in general^16,33,35^, and others exploring specific examples^8,32^. These papers suggest different reasons for the problem, such as the heterogeneous biological variations in some datasets^16,33^ or problems in the methods used that may lead to non-reproducible results^36,37^.

In particular, Zhang *et al*. ^33^ challenge the claim that the non-reproducibility problem in microarray studies is due to poor quality of microarray technology, by showing that inconsistencies occur even between technical replicates of the same dataset. They also show that heterogeneity in cancer pathology would further reduce reproducibility. Ein-Dor *et al*. ^16^ also show the inconsistencies between the results of subsamples of a single dataset, demonstrating that the set of (gene) biomarkers discovered is not unique. They explain that there are many genes correlated with the group outcomes, but the empirical correlations change for different (sub)samples of instances. These two papers motivate our need for tools that can effectively estimate the reproducibility – such as the ones presented here.

Several projects^35–37^ have attempted to formally analyse this problem. Ein-Dor *et al*. ^35^ describe a method, probably approximately correct (PAC) sorting, that estimates the minimum number of instances needed for a desired level of reproducibility. As an example, this worst-case analysis proves that, to guarantee a 50% overlap between different gene lists for breast cancer, each dataset needs to include at least several thousand patients. This suggests poor repeatability results when using small sample sizes, which is consistent with our results for datasets with smaller sample sizes; see Subsection 3, especially Figure 4.

The goal of the MicroArray Quality Control (MAQC) project^36^ was to address the problems and uncertainties about the microarray technology that were caused by the observation that different studies (of the same phenotype) often found very different biomarkers. They suggest that the common approach of using just t-test *p*-values (especially with stringent *p*-values) can lead to poor reproducibility, which motivated them to consider methods like fold-change ranking with a non-stringent *p* cutoff, which they demonstrate leads to more reproducible gene sets. In a follow-up, Guo *et al*. ^37^ found similar results by using the same procedures for another dataset. However, Klebanov *et al*. ^38^ later show that these MAQC project results do not prove that using t-tests is necessarily unsuitable – *i.e*., just because another method (here fold-change) can generate more reproducible results, does not mean that it is performing better; as an extreme, the algorithm that declares every gene is a biomarker (think *p* = 1.0), is completely reproducible. They demonstrate these points by using a set of simulation studies (where they know the “true biomarkers”), and use either t-test or fold-change to propose potential biomarkers. These studies found that the t-test approach performed much better than the fold-change, in terms of recall (sensitivity). These results motivated us to use the t-test approach (rather than fold-change) as our main BD algorithm – which we use for all of our empirical experiments.

Our approximation algorithms use a type of re-sampling to bound reproducibility. Below we summarize several other studies that similarly deal with re-sampling and biomarker discovery. Some studies provide ways to better estimate the true statistical significance, but do not provide a framework for evaluating empirical reproducibility – *e.g*., Gagno *et al*. ^39^ used bootstrap resampling to estimate 95% confidence intervals and *p*-values for an internal assessment of their findings (related to breast cancer survival), Chitpin *et al*. ^40^ proposed a resampling-based method to better estimate the false discovery rate in chromatin immunoprecipitation experiments, and Pavelka *et al*. ^41^ proposed a resampling-based hypothesis testing algorithm that provides a control of the false positive rate for identification of differentially expressed genes. Furthermore, Alshawaqfeh *et al*. ^42^ and Zhao and Li^43^ suggested methods for consistent biomarker detection in high-throughput datasets where evaluation was based on common biomarkers among the two resampled sets – *i.e*., they are considering the false positives but not false negatives. (By contrast, our use of Jaccard score involves both.) Other studies had different goals – *e.g*., Ma *et al*. ^44^ used resampling in a permutation test, to evaluate the predictive power of the identified gene set based on the accuracy of down-stream classification task; recall however that our goal is to evaluate the reproducibility of the biomarkers directly (not downstream). Filosi *et al*. ^45^ propose methods for evaluating the stability of reconstructed biological networks in terms of inference variability due to data subsampling; we however are focusing on the reproducibility of the individual biomarkers. Hua *et al*. ^46^ conducted a simulation study to compare the *ranking performance* of several gene set enrichment methods; by contrast, our approach considers the SET of biomarkers, not the ranking, and is over several real-world datasets (not just simulated ones). Note that none of these used re-sampling techniques to bound the expected replicability of the set of biomarkers found by some discovery algorithm, nor to demonstrate the validity of those bounds.

## 2 Methods

### 2.1 Formal Description

As illustrated in Figure 1, a “Biomarker Discovery” algorithm, BD(·)), takes as input a dataset *D* of *n* subjects, each described by *r* features *F* = {*f*_1_, …, *f*_*r*_} and labeled with a binary outcome, and returns a subset *F*′ ⊂ *F* of purported biomarkers.

Typically, each *f* ∈ *F*′ differs in some significant way between each class. To be more precise, let 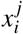 be the value of the *i*^*th*^ feature of the *j*^*th*^ subject, and 𝓁^(*j*)^ be the outcome of the *j*^th^ subject (which is either + or -). Then a class difference means that the set 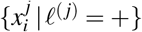 of values of the *i*^th^ feature of the diseased individuals is significantly different from the values of that feature over the healthy individuals, 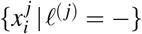. For simplicity, we will assume that these 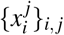 values are either all continuous (such as height, or the expression value of a gene), or all discrete (such as gender, or the genotype of a SNP). Subsection 2.2 below will describe several such biomarker discovery algorithms.

As noted in Equation 1, the *Reproducibility Score* RS(*D*, BD) quantifies the “reproducibility” of the set of proposed biomarkers BD(*D*) corresponding to running the biomarker discovery algorithm over the labelled dataset *D*. Here, we assume that the values of each feature 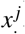, for each outcome *c*, are generated independently from a fixed distribution (*i.e*., “i.i.d.”)

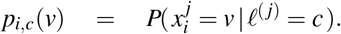

Here and in general, we use *P*(·) to refer to either a probability density for continuous variables, or a probability mass for discrete variables. Note these are just the marginal distributions: we do not assume that the various features are independent from one another. – *i.e*., this does not necessarily correspond to Naive Bayes^30^. We will view 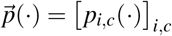 as the matrix of these *r* × 2 different distributions, and let 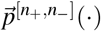 be the distribution for sampling *n* = *n*_+_ + *n*_−_ instances independently from this distribution, where *n*_+_ ∈ ℤ^+^ instances are drawn from the distribution [*p*_1,+_(·), …, *p*_*r*,+_(·)] associated with positive outcomes, and similarly *n*_−_ ∈ ℤ^+^ instances from the distribution [*p*_1,−_(·), …, *p*_*r*,−_(·)] associated with negative outcomes. Then for datasets 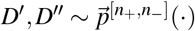 sampled independently, we define

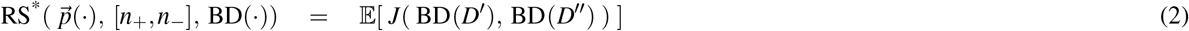

where the Jaccard score of two sets

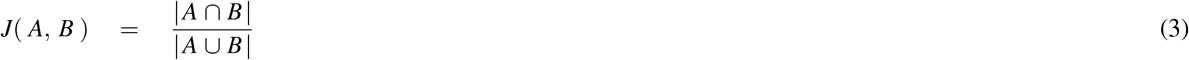

is the ratio of the intersection to the union of these sets – hence *J*(*A, B*) ranges from 0 to 1, and is 1 if and only if *A* = *B* ≠ {}, and is 0 if and only if these sets are disjoint. (We define this to be 0 if *A* = *B* = {}.) Note that the Jaccard score is only one possible measure to evaluate the degree of overlap of gene signatures; Appendix B.4 discusses a slightly different measure that is sometimes used to measure the reproducibility of a set of biomarkers. See also Shi *et al*. ^47^ for a comprehensive overview of these measures.

Of course, we do not know 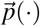, and so we use an empirical distribution 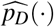, determined based on context from the dataset *D*, to produce the approximation

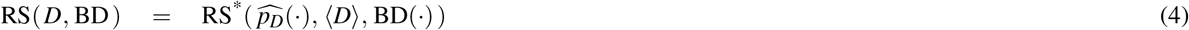

that estimates the reproducibility of the biomarker set BD(*D*), where the notation ⟨*D*⟩ = [|*D*^+^|, |*D*^−^|] refers to the pair of sizes of the positive and negative subjects in *D* – corresponding to [*n*_+_, *n*_−_]. Note that this reproducibility score deals with the *sets* of biomarkers that are produced by the BD(·) function, and not any specific biomarker.

Of course, Equation 4 suggests the obvious bootstrap sampling algorithm ^48^. Empirically, however, we found that it did not perform well (see Appendix B.1) – motivating the algorithms described in Subsection 2.3.

### 2.2 Biomarker Discovery Algorithms: BD(·)

We now discuss various approximation algorithms for biomarker discovery. As our goal is to illustrate the reproducibility issues with respect to the standard approach, we focus on that standard approach: where the biomarker discovery is based on independent two-sample *t*-tests, perhaps with family-wise error correction (see below). Note, however, that the methodology used here is completely general, and can easily extend to any other class of biomarker discovery process; see Limitations in Section 4.

Recall that we considering two types of datasets, depending on whether its feature values (the 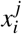 mentioned above) are continuous or discrete. However, for datasets with categorical values – SNPs in our analysis – we use a simple preprocessing step, which precedes all the BD(·) algorithms described here, to convert each categorical value to a real number: here converting each SNP feature, which ranges over the values {AA, Ab, bb}, to the real-values {0, 1, 2}, corresponding to the number of minor alleles (“b”) in the genotype. This allows us to view each such dataset as one with continuous values.

We assume that the real values of each feature, for each outcome, follows a normal distribution, which might be different for the different outcomes, and so we use an independent two-sample *t*-test for all of our empirical experiments. Recall that the test statistic is given by

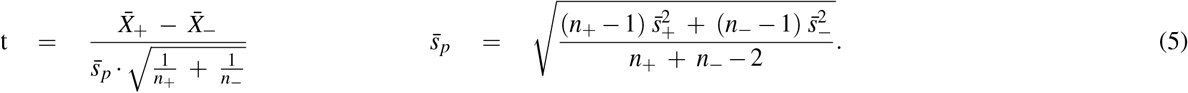

where *n*_+_ and *n*_−_ are the number of instances with positive and negative outcomes, respectively, and with empirical means 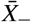 and 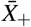 and empirical variances 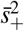 and 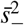.

Note the biomarker discovery process essentially performs a single statistical test for each feature. As the number of features is often large – often tens-of-thousands, or more – many projects sought ways to reduce the chance of false discoveries; a standard way to do this is through some MCC correction. We therefore consider biomarker discovery algorithms described as BD_*t,p,χ*_ (*D*), where the *t* in the subscript refers to the 2-sided *t*-test, the *p* for the *p*-value used, and *χ* to the MCC correction. Our canonical example is BD_*t*,0.05,*BH*_(*D*), with *p* =0.05, and *χ* =BH to the Benjamini/Hochberg correction^49^. This notation makes it easy to consider many variants — *e.g*., adjusting the *p*-value used for the statistical test, whether it is applying another multiple testing correction, or none, etc. See Appendix B.2 for more details.

### 2.3 Algorithms that Approximate the Reproducibility Score

As we have the dataset *D* with *n* ≈ [*n*_+_, *n*_−_] labeled instances, we can directly compute BD(*D*). To compute RS(*D*, BD(·)), however, we also need to produce one (or more) similar datasets *D*′, each with [*n*_+_, *n*_−_] subjects drawn from the same (implicit) distribution 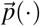 that generated *D* (with the same number of positive and negative instances), but which is presumably disjoint from *D*. While we do not have such *D*′ ‘s, and so cannot directly compute the Reproducibility Score, we show below how to compute an overbound and an underbound of RS(*D*, BD(·)).

#### 2.3.1 Overbound: oRS

The oRS(*D*, BD(·), *k*) procedure produces (an estimate of) an overbound for RS(*D*, BD(·)), by making it *easier* for a feature to be selected to be in both purported biomarker sets. In this algorithm, *D* is our given fixed dataset, BD(·) is the given biomarker discovery algorithm, and *k* is a parameter to determine the number of trials when computing the overbound. (Here, and below, see the Glossary in Appendix A.1 for a summary of the algorithms and their arguments.) The oRS algorithm first defines a size-2*n* dataset *DD* that contains two copies of each subject in *D*, of course with the same outcome both times. It then randomly partitions this *DD* into two disjoint size-*n* datasets *D*_1_ and *D*_2_, balanced by outcome. Here, each partition is with respect to the *list* of elements, so it will include duplicates. To insure that the resulting datasets are balanced, oRS first split *DD* into *DD*^+^ and *DD*^−^, where *DD*^+^ are the cases and *DD*^−^ the controls. It then forms 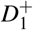 by randomly drawing 1/2 of *DD*^+^, and 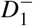 by randomly drawing 1/2 of *DD*^−^, then merges 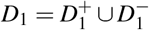; see Figure 2. The dataset *D*_2_ is then formed from the remaining subjects of *DD* not included in *D*_1_. oRS then runs BD(·) on the datasets *D*_1_ (resp., *D*_2_) to produce two sets of biomarkers, and computes the Jaccard score for this pair of biomarker sets: *J*(BD(*D*_1_), BD(*D*_2_)). It then repeats this double-split-BD-Jaccard process *k* times, then returns the average of these *k* values:

**Figure 2.**
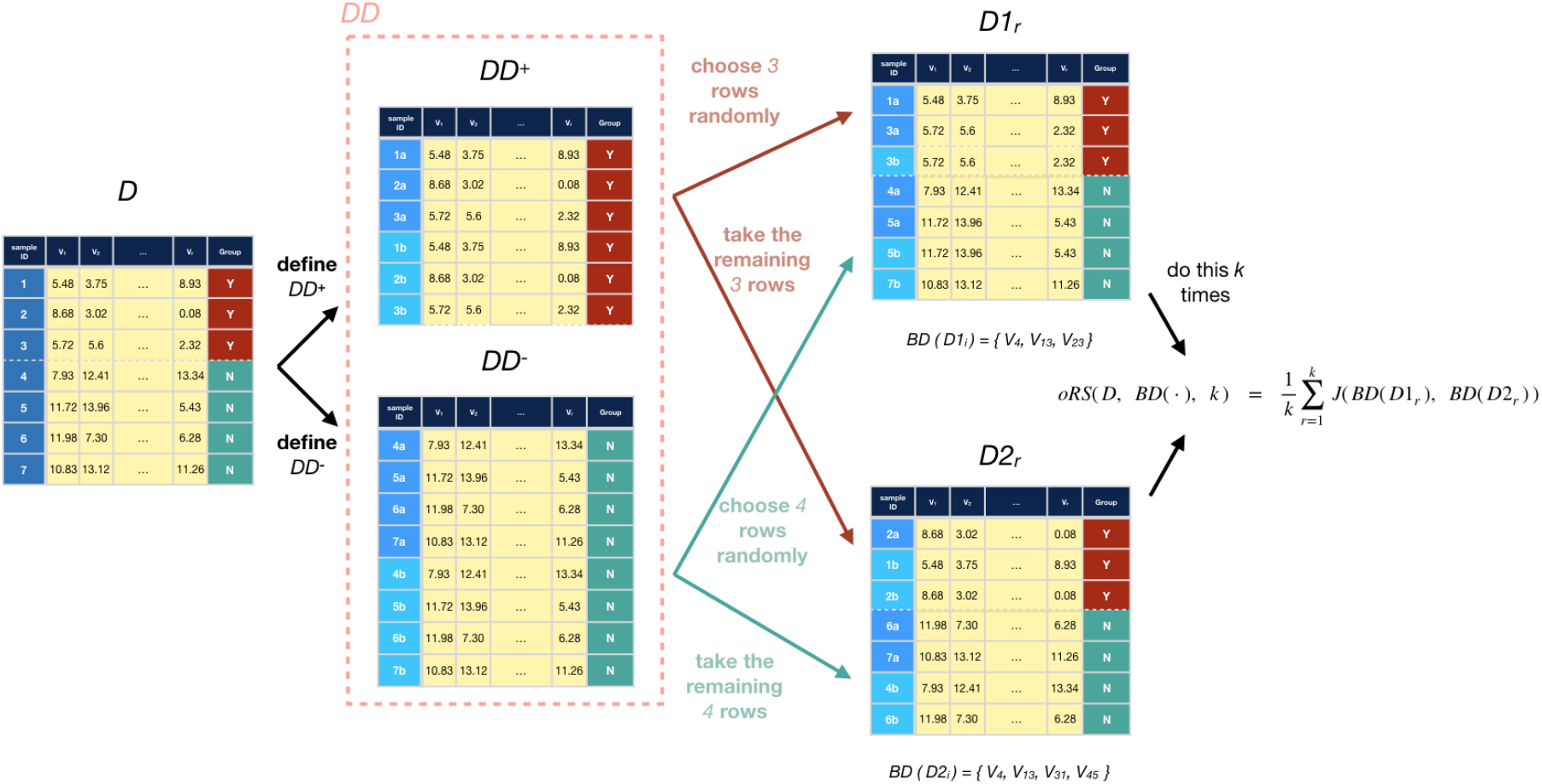
Diagram showing oRS’s process of generating pairs of subsets for a dataset *D* (*k* times), then using those to compute oRS(D, BD(·), k).

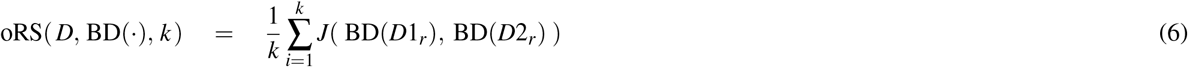

where each dataset pair [*D*1_*r*_, *D*2_*r*_] is created independently using the above procedure.

For each *r*, as we expect *D*1_*r*_ to overlap with *D*2_*r*_, it is relatively likely that any *D*1_*r*_-biomarker will also be a *D*2_*r*_-biomarker (more likely than if *D*1_*r*_ was disjoint from *D*2_*r*_), which means we expect the associated Jaccard score to be higher. This follows from the heuristic that, as two datasets have more common elements, we expect the number of biomarkers common to two datasets, to increase – *i.e*.,

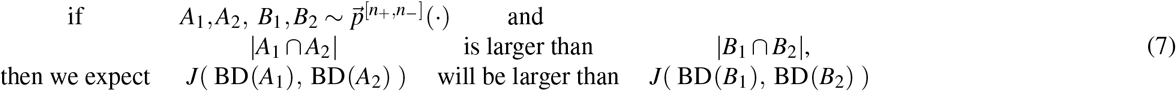

*ceteris paribus*. Here, we view each of {*A*_1_, *A*_2_, *B*_1_, *B*_2_}as a set of *n* = *n*_+_ + *n*_−_ *r*-dimensional instances. Note this relationship is simply a heuristic to motivate the algorithm – one that we expect to hold in practice. Appendix C provides some basic arguments, and empirical evidence, to support this claim.

We close with three quick observations:

1. **Expected Overlap:** We expect 50% of the instances to be duplicated in any given pair of datasets. See Lemma 2 in Appendix C.4.
2. **Relation to Bootstrap Samples:** Here, each subject occurs exactly twice in each pair of datasets *D*1_*r*_ and *D*2_,*r*_. If we instead used bootstrap sampling (called bRS below), we expect many subjects would occur more often in the pair of datasets. (See Lemma 3 in Appendix C.4.) Given Heuristic 7, this means we expect bRS’s Jaccard score here to be higher than for oRS’s; as oRS is already an overbound for the true score (RS), this means bRS would be a worse bound, as bRS ≥ oRS ≥ RS. This is why we use our “doubling approach” oRS rather than bootstrap sampling bRS, as oRS produces values that are smaller, but still remains an overbound, as desired. See Figures 7 and 8 and Appendix B.1.
3. **Relation to** RS^*^: Our oRS approach is clearly related to RS^*^ (Equation 2), as both compute the average Jaccard score of pairs of size-*n* datasets sampled from a distribution. They differ as (a) RS^*^ uses the true distribution 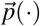, while oRS uses only the estimate 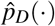, (b) RS^*^ is the *true* average while oRS is just the *empirical* average over *k* trials, and (c) RS^*^ will draw *independent* datasets, but the oRS datasets will overlap.

#### 2.3.2 Underbound: uRS

The uRS(*D*, BD, *k*) procedure produces (an estimate of) an underbound for RS(*D*, BD(·)) by making it *harder* for a feature to be selected to be in both purported biomarker sets. First, observe that as [*n*_+_, *n*_−_] increases (keeping the *n*_+_-to-*n*_−_ ratio fixed, as we consider changing the size of the dataset), we expect the statistical estimates to be more accurate, and in particular, statistical tests for differences between the two classes will be correct more often. Hence, a statistical test will better identify the “true” biomarkers *F*^∗^ from a size-*n* subset *D*^(*n*)^, versus from a size-*n/*2 subset *D*^(*n/*2)^. Now consider two size-*n* datasets 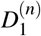 and 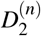, and also two size-*n/*2 datasets 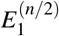 and 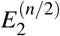. As 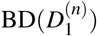 and 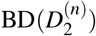 are each closer to *F*^∗^ than 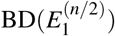 and 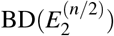, we expect 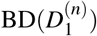 and 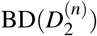 to be closer to each other, than 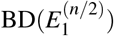 and 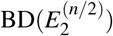, which means we expect that

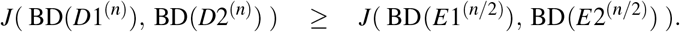

In general, given that

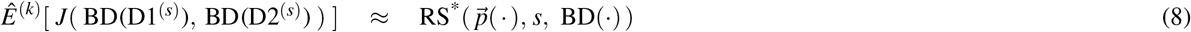

(where *Ê*^(*k*)^[·] is the empirical average over *k* samples), this argues that the RS^*^ score should increases with the size *s* of the dataset – which suggests that

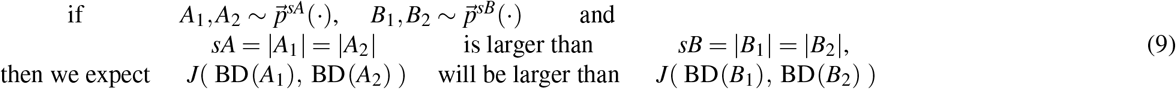

Figure 4(a) presents empirical evidence, over 5 datasets, supporting this claim – showing that the Jaccard score increases as we increase the size *s* of the datasets. Appendix C provides some arguments, and additional empirical evidence (over hundreds of simulations), that further support this heuristic.

This motivates our underbound algorithm uRS(*D*, BD, *k*), which first partitions *D* into two disjoint size-*n/*2 subsets, *E*_1_ and *E*_2_, with balanced outcomes. It then computes *J*(BD(*E*_1_), BD(*E*_2_)) which, assuming Heuristic 9, is an underbound in expectation for RS(*D*, BD(·)). uRS does this partitioning *k* times, producing *k* different dataset pairs {[*E*_1,*r*_, *E*_2,*r*_]}_*r*=1,…,*k*_, and returning the average Jaccard score, i.e.

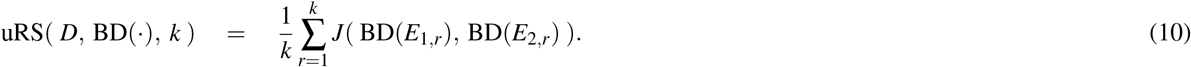

See Figure 3.

**Figure 3.**
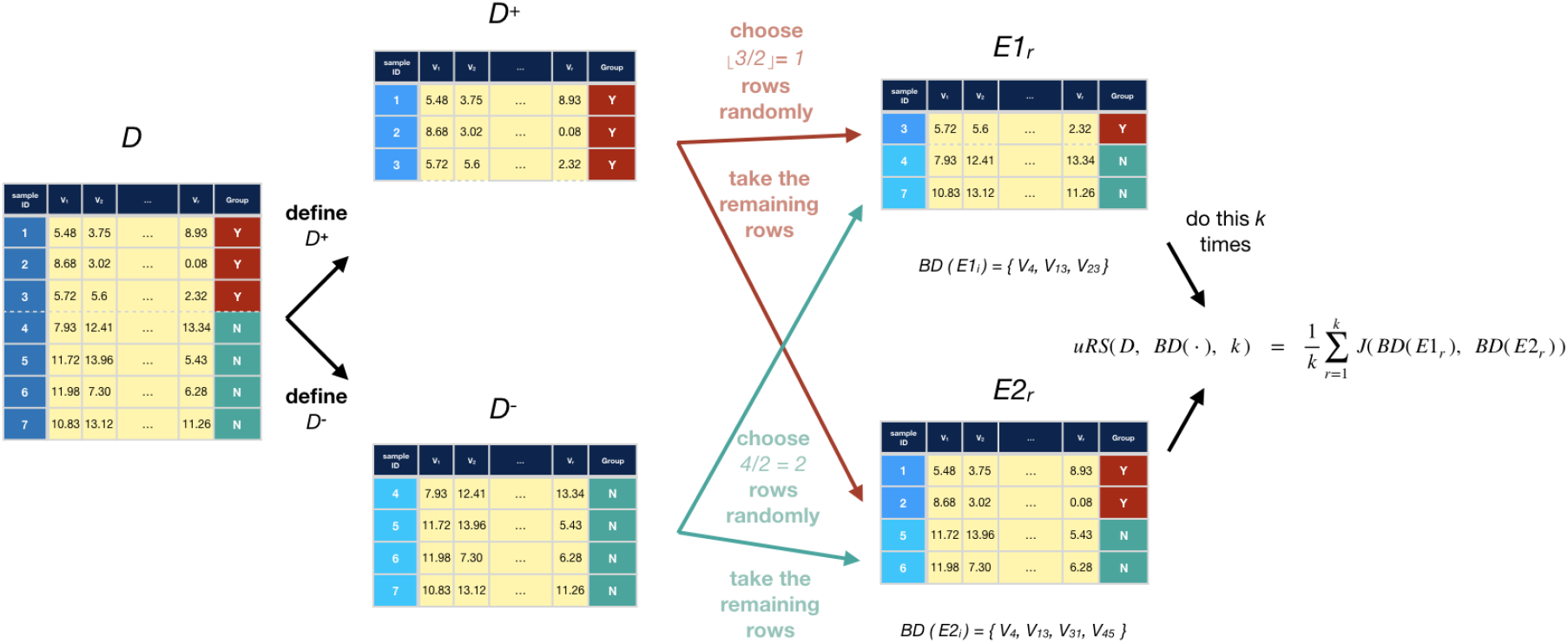
Diagram showing uRS’s process of generating pairs of subsets for a dataset *D* (*k* times), then using those to compute uRS(D, BD(·), k).

**Figure 4.**
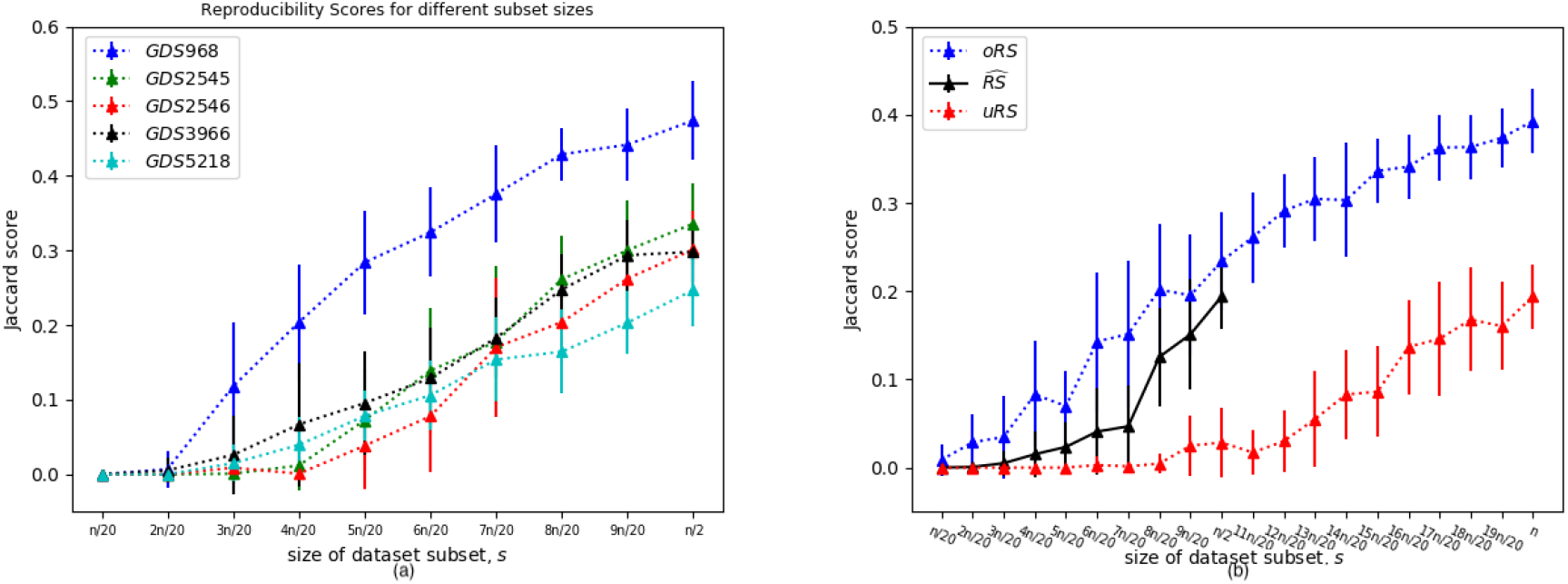
(a) For each of the 5 datasets *D*, each point shows the average (±sd) Jaccard *Ê*^(*k*)^[*J*(BD(*D*1^(*s*)^), BD(*D*2^(*s*)^))] over *k* = 20 pairs [*D*1^(*s*)^, *D*2^(*s*)^] of disjoint size-*s* subsets of *D*. Here, *n* is the size of the original dataset – note this can only go to *n/*2 – and we are using the standard BD_*t*,0.05,*BH*_. (b) Showing how the approximations relate to one another, and scale with the size *s* of the dataset. Here we are using subsets of the Metabric dataset, with *n*=1654. We observed the same behavior for all datasets.

### 2.4 Empirical study over Various Datasets

There are now many publicly-available datasets that have been used in association studies. Here, we use them to …

U1: Better understand what Jaccard scores are typical, for the standard BD(·) algorithms;

U2: Determine whether our predictions match the results of earlier meta-analyses; and

U3: Determine if our approximations are meaningful – *i.e*., if (for reasonable values of *k*):

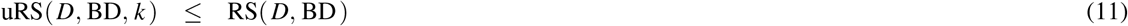

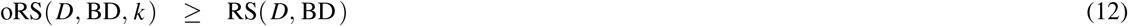

The next section will explicitly discuss all three issues. Of course, given only a single dataset *D* of size-*n*, we cannot compute, nor even estimate, the true value of RS(*D*, BD(·)). However, we can estimate RS(*D*^(*n/*2)^, BD(·)), where *D*^(*n/*2)^ is a size-*n/*2 (outcome-balanced) subset of *D*. In fact, uRS(*D*, BD(·), k) is a meaningful estimate of RS(*D*^(*n/*2)^, BD(·)); below we will use

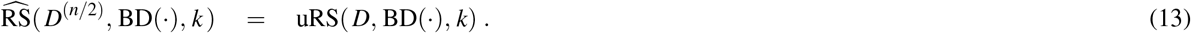

We will then compare this 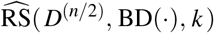 against uRS(*D*^(*n/*2)^, BD, *k*) and oRS(*D*^(*n/*2)^, BD, *k*) and to see whether the relations of Equations 11 and 12 both hold, with respect to various size-*n/*2 subsets *D*^(*n/*2)^.

More generally, we can do this for any size-*s* subset *D*^(*s*)^ of *D* where *s* ≤ *n/*2. Here, we need a set of pairs of disjoint outcome-balanced subsets *D*′, *D*′′ ⊂ *D* where |*D*′ | = |*D*′′ | = *s* and *D*′ ∩ *D*′′ = {}. For a fixed dataset *D*, and specified number *k* ∈ ℤ^*>*0^, we can then plot these 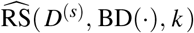, BD(·), *k*) values along with oRS(*D*^(*s*)^, BD, *k*) and uRS(*D*^(*s*)^, BD, *k*), as a function of *s* to see their behaviour; see Figure 4(b), for the Metabric dataset. Our website https://biomarker.shinyapps.io/BiomarkerReprod/ also provides this visualization.

We explored our approximations over **25 different real-world datasets, including 16 microarray datasets and 2 RNAseq datasets with continuous data, and 7 SNP datasets with categorical data** (see Table 1). This first set includes 4 of the gene expression datasets discussed in the Zou *et al*. ^8^ meta-analysis – each describing metastatic versus non-metastatic breast primary cancer subjects – to see if our method is consistent with their empirical results. We also included 11 other relatively-small gene expression datasets (with 19 to 187 subjects), focusing on human studies that had a binary class outcome from the GEO repository. To explore how our tools scale with size, we also included 3 other relatively large datasets, with 532 to 1654 subjects. As these were survival datasets, we set the binary outcome based on the median survival time (removing any subject that was censored before that median time). In addition to these 4+11+3 = 18 gene expression datasets (with real-valued entries), we also include 7 SNP datasets (with 39 to 164 subjects), with discrete values, also selected from human studies with binary class outcome s. Figure 5 plots the number of features and biomarkers found, using the BD_*t*,0.05,*BH*_ algorithm, for each dataset – both *D*^(*n*)^ and *D*^(*n/*2)^.

**Table 1.**
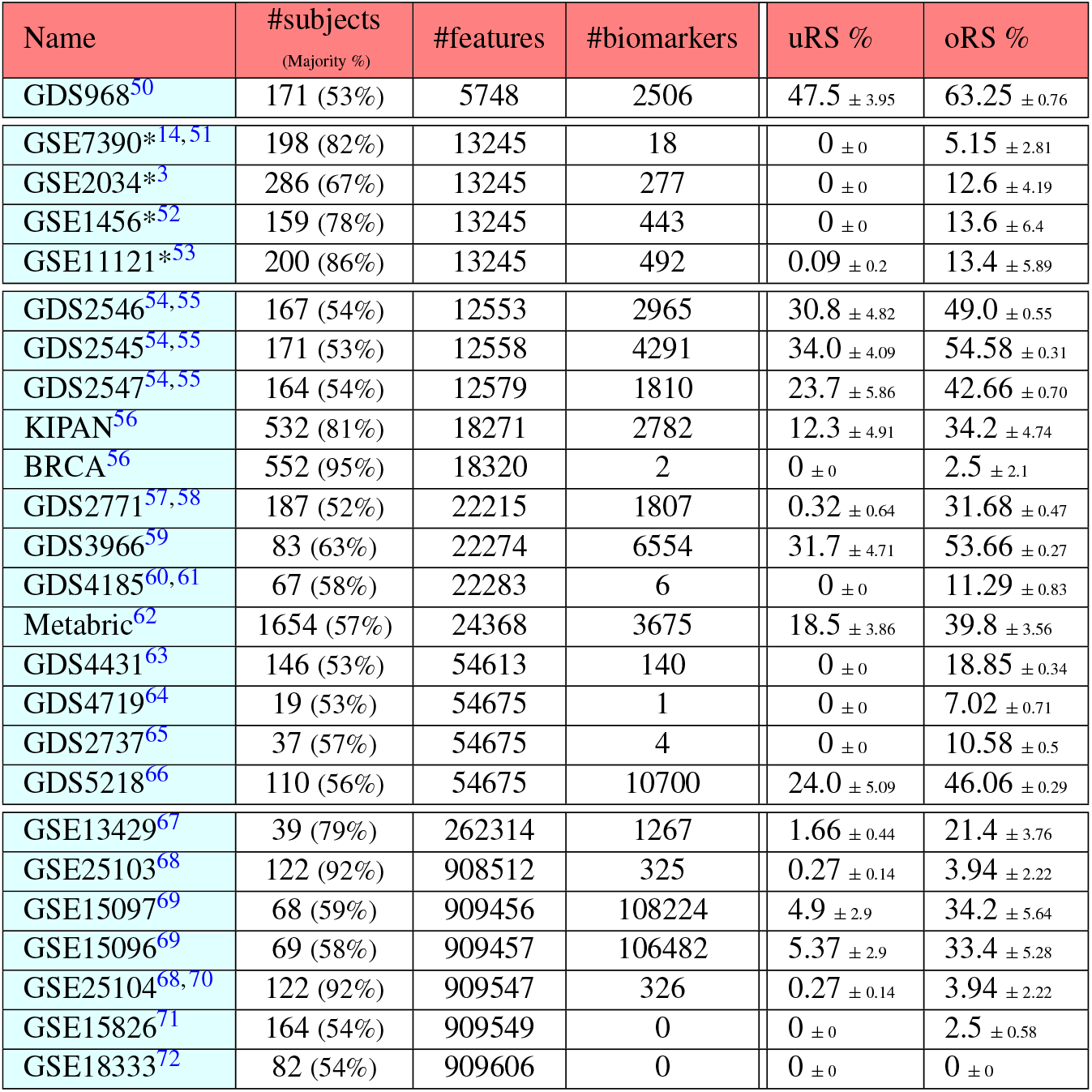
Results for all 25 datasets when using all the subjects, sorted by sample size (#subjects) – corresponding to Figure 5. The first 18 entries are Gene Expression datasets (including the 4 “*”ed entries, from the Zou *et al*. ^8^ meta-study), and the final 7 are SNP datasets. Reproducibility Scores are shown in the form of mean ± standard deviation. The “(Majority %)” values are the percentage of the subjects in the dataset with the more common outcome – *e.g*., 53% of the subjects in the GDS968 dataset are labeled “+” (for “long survival time”), and 82% of GSE7390 are labeled “−” (for “Non-Metastatic”).

| Name | #subjects<br>(Majority %) | #features | #biomarkers | uRS % | oRS % |
| --- | --- | --- | --- | --- | --- |
| GDS968<sup>50</sup> | 171 (53%) | 5748 | 2506 | 47.5 ± 3.95 | 63.25 ± 0.76 |
| GSE7390*<sup>14,51</sup> | 198 (82%) | 13245 | 18 | 0 ± 0 | 5.15 ± 2.81 |
| GSE2034*<sup>3</sup> | 286 (67%) | 13245 | 277 | 0 ± 0 | 12.6 ± 4.19 |
| GSE1456*<sup>52</sup> | 159 (78%) | 13245 | 443 | 0 ± 0 | 13.6 ± 6.4 |
| GSE11121*<sup>53</sup> | 200 (86%) | 13245 | 492 | 0.09 ± 0.2 | 13.4 ± 5.89 |
| GDS2546<sup>54,55</sup> | 167 (54%) | 12553 | 2965 | 30.8 ± 4.82 | 49.0 ± 0.55 |
| GDS2545<sup>54,55</sup> | 171 (53%) | 12558 | 4291 | 34.0 ± 4.09 | 54.58 ± 0.31 |
| GDS2547<sup>54,55</sup> | 164 (54%) | 12579 | 1810 | 23.7 ± 5.86 | 42.66 ± 0.70 |
| KIPAN<sup>56</sup> | 532 (81%) | 18271 | 2782 | 12.3 ± 4.91 | 34.2 ± 4.74 |
| BRCA<sup>56</sup> | 552 (95%) | 18320 | 2 | 0 ± 0 | 2.5 ± 2.1 |
| GDS2771<sup>57,58</sup> | 187 (52%) | 22215 | 1807 | 0.32 ± 0.64 | 31.68 ± 0.47 |
| GDS3966<sup>59</sup> | 83 (63%) | 22274 | 6554 | 31.7 ± 4.71 | 53.66 ± 0.27 |
| GDS4185<sup>60,61</sup> | 67 (58%) | 22283 | 6 | 0 ± 0 | 11.29 ± 0.83 |
| Metabric<sup>62</sup> | 1654 (57%) | 24368 | 3675 | 18.5 ± 3.86 | 39.8 ± 3.56 |
| GDS4431<sup>63</sup> | 146 (53%) | 54613 | 140 | 0 ± 0 | 18.85 ± 0.34 |
| GDS4719<sup>64</sup> | 19 (53%) | 54675 | 1 | 0 ± 0 | 7.02 ± 0.71 |
| GDS2737<sup>65</sup> | 37 (57%) | 54675 | 4 | 0 ± 0 | 10.58 ± 0.5 |
| GDS5218<sup>66</sup> | 110 (56%) | 54675 | 10700 | 24.0 ± 5.09 | 46.06 ± 0.29 |
| GSE13429<sup>67</sup> | 39 (79%) | 262314 | 1267 | 1.66 ± 0.44 | 21.4 ± 3.76 |
| GSE25103<sup>68</sup> | 122 (92%) | 908512 | 325 | 0.27 ± 0.14 | 3.94 ± 2.22 |
| GSE15097<sup>69</sup> | 68 (59%) | 909456 | 108224 | 4.9 ± 2.9 | 34.2 ± 5.64 |
| GSE15096<sup>69</sup> | 69 (58%) | 909457 | 106482 | 5.37 ± 2.9 | 33.4 ± 5.28 |
| GSE25104<sup>68,70</sup> | 122 (92%) | 909547 | 326 | 0.27 ± 0.14 | 3.94 ± 2.22 |
| GSE15826<sup>71</sup> | 164 (54%) | 909549 | 0 | 0 ± 0 | 2.5 ± 0.58 |
| GSE18333<sup>72</sup> | 82 (54%) | 909606 | 0 | 0 ± 0 | 0 ± 0 |

**Figure 5.**
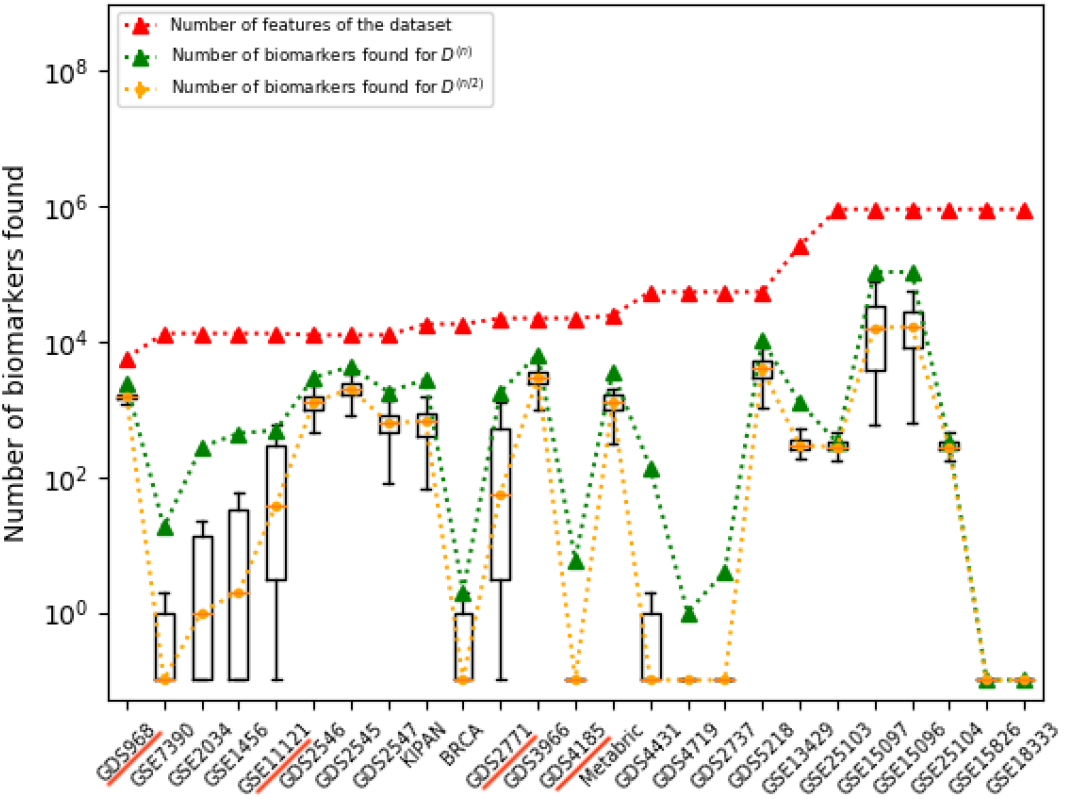
Box+whiskers plots showing number of biomarkers found for *D*^(*n/*2)^ when using BD_*t*,0.05,*BH*_ over *k* = 20 iterations for various dataset, compared to the number of biomarkers for *D*^(*n*)^, and to the number of features in each dataset. Note the y-axis is a log-scale. (Note we first changed all “0” values to “10^−1^”.) For details, see Tables 1 and 2.

## 3 Results

We ran our suite of methods over the aforementioned 25 datasets, including 16 microarray datasets and 2 mRNAseq datasets, whose feature-values 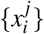 (recall each 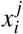 is the expression value of the *i*-th gene for the *j*-th subject; we log_2_-transformed the values from the mRNAseq datasets) and 7 were SNP datasets with categorical entries, *i.e*., 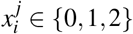 is the number of minor alleles in the genotype for the *i*^th^ SNP for the *j*^th^ subject; see Table 1. Here, we use the standard BD_*t*,0.05,*BH*_(·) biomarker discovery algorithm; see Section 2.2.

**Table 2.**
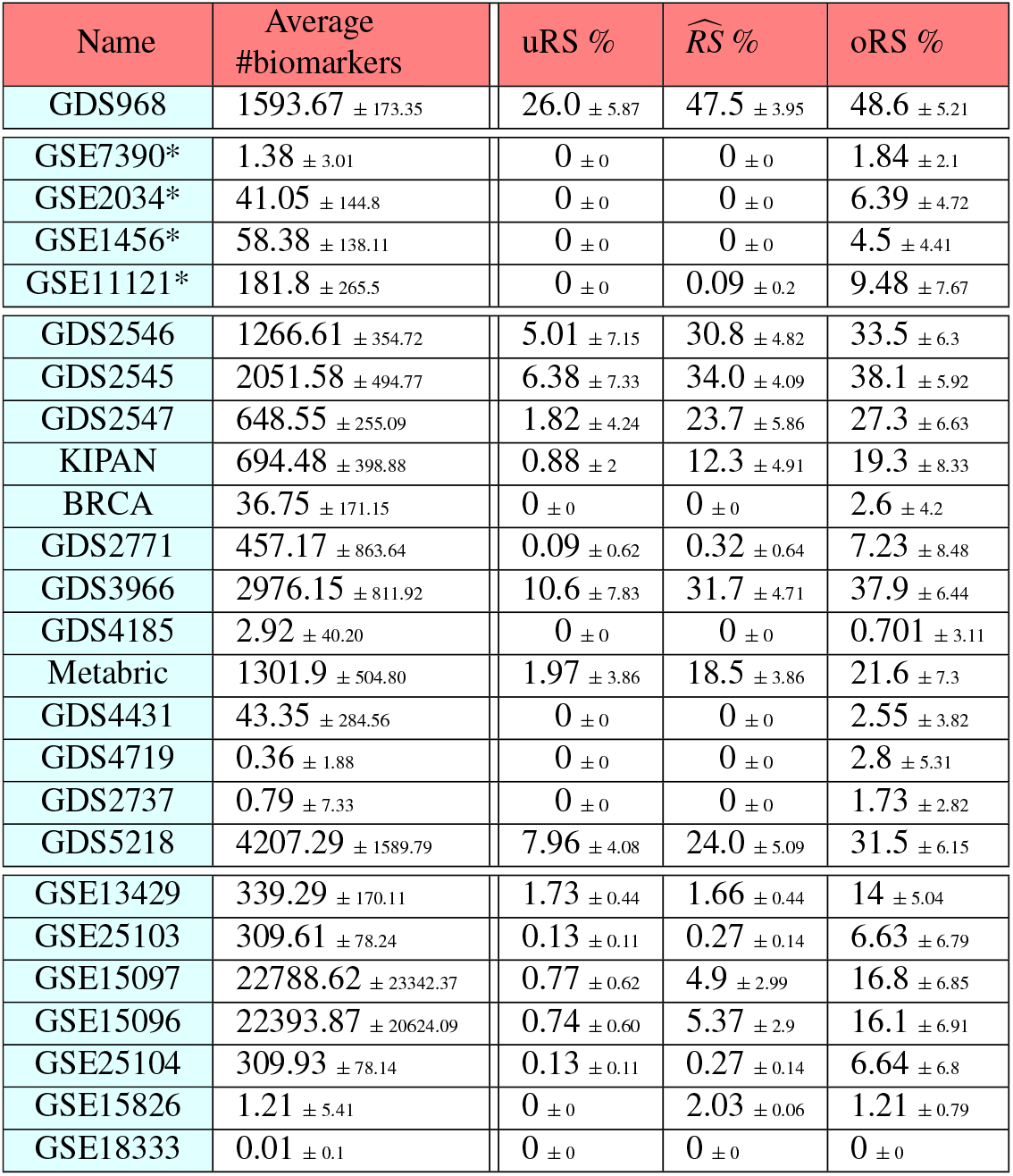
Results for all datasets when using half of the subjects – *i.e*., *D*^(*n/*2)^. Reproducibility Scores and average number of biomarkers are shown in the form of mean ± standard deviation. (The caption for Table 1 describes the row ordering.)

First, to address (U1) and (U2) in Section 2.4, we analyzed the 4 datasets mentioned in the Zou *et al*. ^8^ meta-analysis (see the 4 “*” rows of Table 1) and computed the {uRS(*D*, BD_*t*,0.05,*BH*_, 50), oRS(*D*, BD_*t*,0.05,*BH*_, 50)} values for each dataset *D*, as well as the actual Jaccard score for biomarkers for each pair of datasets. (We were not able to replicate the results reported for the 5^th^ dataset from that study, and so we excluded that one dataset from our analysis.) The results, in Figure 6, show that the Jaccard score for each pair is well within the bounds computed by our approximations, for each of the datasets in that pair – that is, the results for 4 × 3 = 12 ordered-pairs of datasets are consistent with our predictions. Note that we verified that our BD_*t*,0.05,*BH*_ algorithm matched the original study by verifying that the PO scores (Equation 15 in Appendix B.4) matched the ones that were originally published.

**Figure 6.**
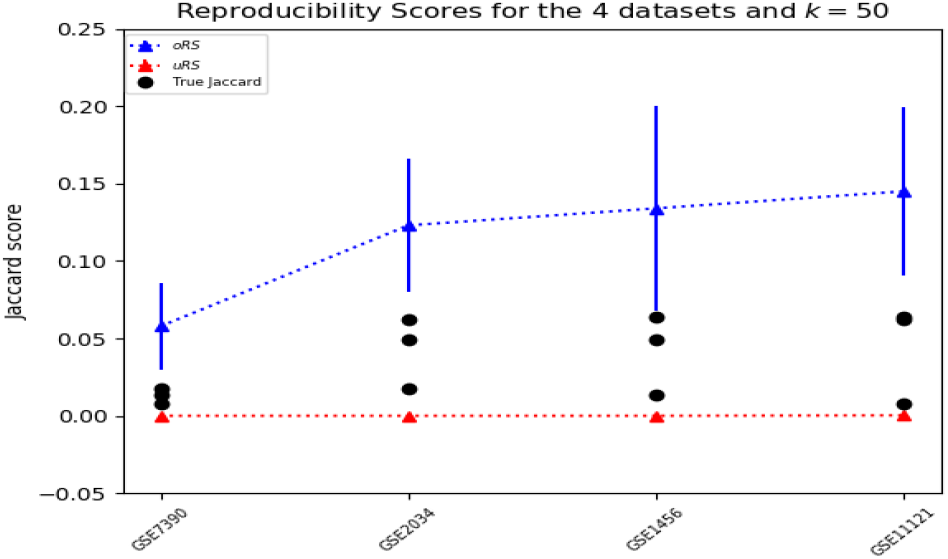
Under-bound and over-bound for the 4 datasets from Zou *et al*. ^8^, as well as the true Jaccard score for each pair – 3 numbers for each dataset, shown by black circles.

**Figure 7.**
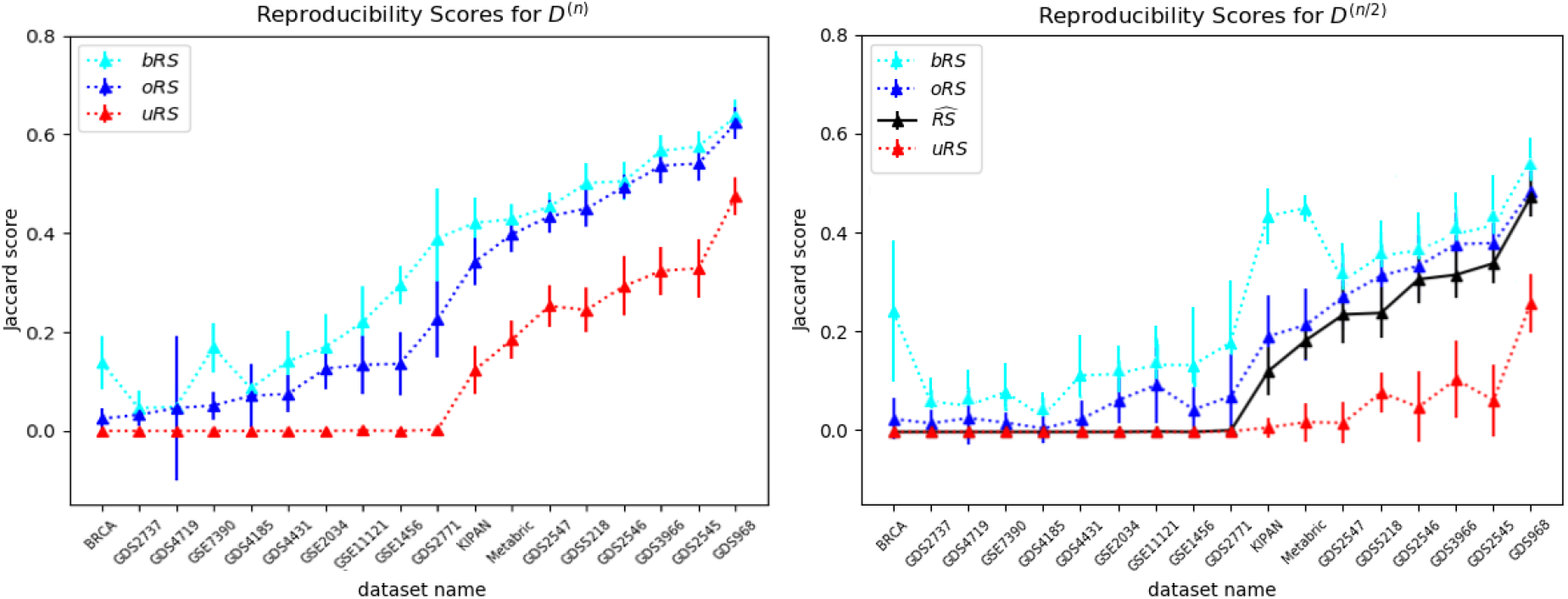
Reproducibility scores (mean and standard deviation) for all 18 continuous datasets, both for complete datasets with *n* subjects (left) and for half-sized with 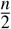 subjects (right), for *k* = 50 iterations. The x-axes (for both plots) are sorted by the value of the over-bound for the *D*^(*n*)^ datasets. We see, in both, that the over-bound oRS is consistently higher than the under-bound uRS. Moreover, the right plot shows that the “truth” 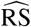 is also between uRS and oRS. (The bRS lines in the plots are based on the Bootstrap Overbound method; see Appendix B.1.)

**Figure 8.**
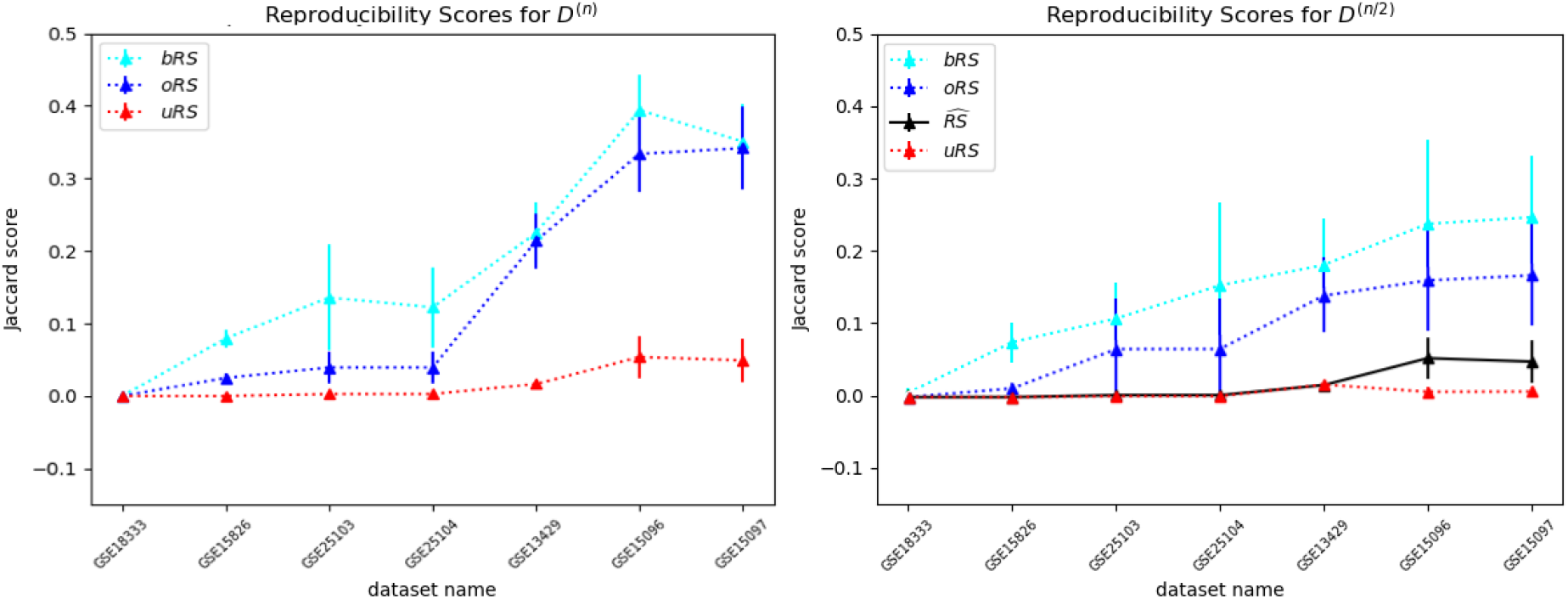
Reproducibility scores (mean and standard deviation) for 7 SNP datasets, both for complete datasets with *n* subjects *D*^(*n*)^ (left) and for half-sized with 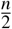 subjects *D*^(*n/*2)^ (right), for 50 iterations. The x-axes (for both plots) is sorted by the value of the overbound oRS for the *D*^(*n*)^ datasets. We see, in both, that the over-bound oRS is consistently higher than the under-bound uRS. Moreover, the right plot shows that the “truth” 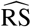 is also within the range of oRS and uRS. (The bRS lines in the plots are based on the Bootstrap Overbound method; see Appendix B.1.)

To address (U3), we also analyzed the other 14 continuous datasets *D*, and computed the oRS and uRS values using *k* = 50 repetitions; see Figure 7[left]. We see that the overbound oRS is consistently larger than the underbound uRS – *i.e*., uRS ≤ oRS – as claimed by Equations 11-12. Figure 7[right] plots the corresponding values for the *D*^(*n/*2)^ datasets, that use only 1/2 of the dataset, using the same BD(·) algorithm and *k* = 50. It also plots the “true” 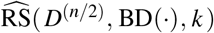 values for the datasets. Again, we see that 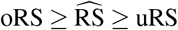.

Finally, similar to that experiment over the 18 continuous datasets, we examined the 7 discrete datasets and produced the reproducibility scores. Figure 8[left] shows the scores for each of the 7 SNP datasets, demonstrating that oRS ≥ uRS holds for the discrete cases as well. Figure 8[right] shows the scores for *D*^(*n/*2)^ datasets, and performs the same verification. Those figures also allow us to see the Jaccard scores (U1 above) range from essentially 0 to around 0.475 for the *D*^(*n/*2)^ datasets. In addition to the plots, Tables 1 and 2 present the relevant values.

Appendix B provides the results of many additional empirical studies, showing how the reproduciblity scores change based on which (if any) MCC correction is used, the specific *p*-value used for the t-test, the number of draws *k* used by the various approximations, and the size of the dataset *n*.

## 4 Discussion

### Limitations

This paper provides an effective way to estimate the reproducibility of the biomarkers found from a dataset, using a biomarker discovery algorithm. While the message of this paper is very general, the specific analyzes here all used the standard discovery algorithm BD_*t*,0.05,*BH*_, to illustrate that the issues apply to the approaches commonly used. Appendix B explores some other discovery methods, that are based on t-tests. We anticipate the same issues would hold for other tests, such as Mann-Whitney, Wilcoxon or even multivariate analysis – but this is future work. All of our empirical studies dealt with standard datasets, whose values were either all real values or all categorical values; none had some of each. We only considered datasets whose outcomes are binary and our use of t-test implicitly assumes the feature values, given each outcome, are Gaussian. Our analytic model considers the overlap of biomarkers found from two datasets, of the same size. (That is, we do not consider how the biomarkers obtained from a size-100 dataset, overlap with those from a size-300 dataset.) Finally, our analysis estimated the expected RS, for a given BD and dataset *D*. It would be interesting to explore a variant of this: given a dataset *D* and a minimum score *s >* 0, find the “BD (*D, s*) discovery algorithm” that would produce the biomarker set whose expected Jaccard score would be at least *s*. (This might mean adjusting the *p*-value cut-off, and/or including some specific MCC algorithm, or some other modification.)

### Contributions

This paper has several contributions: (1) It motivates, then provides, a formal definition of reproducibility, that can help researchers “evaluate” a set of purported biomarkers; (2) It provides a pair of algorithms that can accurately bound this “reproducibility score” for a given dataset and biomarker discovery algorithm; (3) It provides empirical evaluation of these algorithms, over 25 different real-world datasets, to demonstrate that they work effectively; and (4) It introduces a freely-available website https://biomarker.shinyapps.io/BiomarkerReprod/ that runs these algorithms on the dataset entered by a user, and biomarker discovery system, which will allow users to easily evaluate the quality of the biomarker set produced. Given how easy it is to use this tool, we hope that future researchers will automatically use it to quickly evaluate the quality of the purported biomarkers, then include these estimates when they publish their biomarkers. We anticipate this analysis may also lead to new biomarker discovery algorithms, to optimize this reproducibility score.

## Acknowledgements

The authors all gratefully acknowledge the funding support from Canada’s NSERC (Natural Sciences and Engineer Research Council) and the Alberta Machine Intelligence Institute (Amii). We also benefited from the preliminary research work by Patrick Schwaferts and discussions with Sambasivarao Damaraju.

## Author contributions statement

All 4 authors helped design the experiment, analyze the results, and write this manuscript. In addition, R.G. motivated and initiated this project; S.K. designed and implemented the overbound and underbound algorithms, and the web application; and A.F. and A.R. (re)implemented and ran the bulk of the experiments. All authors reviewed and approved the final manuscript.

## A Notes from Text

This appendix provides various short notes related to material in the main text:

### A.1 Glossary

| Term | Description |
| --- | --- |
| $BD_{t,p,\chi}(D)$ | **Biomarker Discovery** algorithm: given a labeled dataset $D$, return the set of features (associated with $D$), that exhibit a class difference; see Equation 1. Here, the $t$ in the subscript refers to the 2-sided $t$-test, the $p$ for the $p$-value used, and $\chi$ to the MCC correction. Our canonical example is $BD_{t,0.05,BH}(D)$, with $p = 0.05$, and $\chi = BH$ corresponding to the Benjamini/Hochberg correction. We use $BD(D)$ for a generic algorithm. |
| biomarker | “a characteristic (feature) that is objectively measured and evaluated as an indicator of normal biological processes, pathogenic processes, or pharmacologic responses to a therapeutic intervention”<sup>1</sup> |
| $D, D^{(n/2)}$ | $D$ refers to a labeled dataset, with $n$ subjects, each described by $r$ features, and an outcome. $D^{(n/2)}$ refers to a dataset formed containing half of the elements of the original dataset $D$ – this is balanced, containing $n_+/2$ +-labeled subjects and $n_-/2$ -labeled subject |
| feature | sometimes called the “independent variables” or “covariates” – here, we assume they are all real valued |
| gene expression | continuous value that indicates how much RNA of a specific gene has been expressed |
| GWAS | genome-wide association study |
| $J(A, B)$ | Jaccard score of pair of sets $A$ and $B$; see Equation 3 |
| $n, [n_+, n_-]$ | By convention, $n$ is the number of subjects in the dataset, which includes $n_+$ subjects that are labeled +, and $n_-$ subjects labeled -. |
| $oRS(D, BD(\cdot), k)$ | (an estimate of) an overbound of $RS(D, BD(\cdot))$ where $k = \#repetitions$; see Equation 6 |
| outcome | sometimes called the “label”, “response” or “dependent variable” – here, just considering binary (think “case vs control”) |
| $\vec{p}(\cdot) \quad \vec{p}^{[n_+, n_-]}(\cdot)$ | $\vec{p}(\cdot) = [p_{j,c}(\cdot)]_{j,c}$ is the $2 \times r$ matrix of the distributions for the 2 outcomes and $r$ features; $\vec{p}^{[n_+, n_-]}(\cdot)$ is the distribution that draws, for each feature $j$, $n_+$ subjects from $p_{j,+}$ and $n_-$ from $p_{j,-}$ |
| $\widehat{p}_D(\cdot)$ | empirical estimate of $\vec{p}(\cdot)$, based on the context of the dataset $D$ |
| $RS(D, BD(\cdot))$ | “Reproducibility Score” of the biomarkers produced, by running biomarker-discovery algorithm $BD(\cdot)$ on the labeled dataset $D$; see Equation 4 |
| $RS^*(\vec{p}(\cdot), [n_+, n_-], BD(\cdot))$ | reproducibility score for the biomarkers produced by the $BD(\cdot)$ biomarker discovery tool on two datasets of size $[n_+, n_-]$ drawn from $\vec{p}(\cdot)$; see Equation 2 |
| SNP | single nucleotide polymorphism – discrete, with values {AA, Ab, bb} indicating homozygous dominant, heterozygous, and homozygous minor, respectively |
| subject | a “record” in the dataset (e.g., a row in the table shown in Figure 1). Also called “element” or “instance” |
| $uRS(D, BD(\cdot), k)$ | (an estimate of) an underbound of $RS(D, BD(\cdot))$, where $k = \#repetitions$; see Equation 10 |

### A.2 How biomarker discovery differs from supervised machine learning

We earlier noted a few differences between association studies versus predictive studies. Here, we list two others:

a. In general, a predictive model provides some information about *an individual* (corresponding to a **row** of the matrix at the top of Figure 1) – *e.g*., whether she has some specific disease. By contrast, an association study identifies *features* (each corresponding to a **column**), with the prediction that each considered a biomarker will each exhibit some *population difference with respect to a dataset of many individuals*.
b. It is relatively easy to evaluate the quality of a learned predictive model, by running that predictor on a held-out set of subjects. By contrast, there is no direct way to determine if a purported biomarker is correct. This is why we, instead, look for “consistency” of a set of biomarker discovery tools. That is, we hope that these discovered feature sets have low variance. (Note that they can have high-bias – *e.g*., if they all set *p* = 1, then each discoverer will return all features; this will have low variance, but presumably high bias.)

### A.3 Why do many papers not provide biological validation?

This may be because such biological validation is not yet implementable, or the technology is not yet available. Alternatively, the biological validation may be possible but be a major project that those authors hope to explore in future works.

### A.4 Our model (Equation 1) deals with just a single step

Some modern GWAStudies involve many phases – typically using one phase to reduce ≈10^6^ features to a few thousand based on one dataset, and then using a second dataset to reduce those features to a sub-subset, etc^73^. Here, our analysis is relevant to any one of these phases; see Figure 1. Also, some studies regress out covariates before finding biomarkers; here, we assume that this has happened and our analysis uses those already-regressed-out values.

### A.5 Some features may only be important in combination

Sometimes a feature may be completely independent of the outcome, by itself, but become relevant, in combination with another (independent) feature. As an example, consider babies *in utero*, whose descriptions each include the Rh blood type of its mother, MRh ∈ {+, −}, and also of its father FRh ∈{+, −}, among other features, where the outcome is the health *H* of that baby. Note that *H* may be completely uncorrelated with MRh, and also completely uncorrelated with FRh – which means neither MRh nor FRh could be a biomarker. However, suppose finding these blood factors are different MHr≠FHr, increases the baby’s risk. Assuming balanced sampling (with an equal number of MRh=+ and MRh= − subjects, and similarly for FRh), this means an effective predictive model would need to include both features, even though neither is a biomarker. Moreover, we typically assume that a feature either increases the risk of a disease in all situations, or always decreases that risk. This in-utero-baby example shows this is not always the case: We see that MRh=+ can sometimes increase the risk (when FRh= −), and other times, decrease the risk (when FRh=+). Hence, a simple linear combination of feature values might not always be appropriate.

While this is an extreme situation – where each feature is completely irrelevant by itself – it is relatively common for a disease to be associated with many minor features; here again, it is possible that none of the features, by itself, shows sufficient class distinction

This also happens when the class is inherently heterogeneous – *e.g*., “headache” can be based on various phenomena, including ischemic stroke, dehydration, migraine, etc., each with various different factors. This is believed to happen with essentially all complex genetic disorders, especially when underlying pathologies are not known.

These situations argue that a *panel* of features can sometimes be more appropriate than individual features. If the model starts with a pre-defined combination of a set of features – *e.g*., a simple average of a specific set of gene expression values, or a set of heterozygous settings in a specific set of SNPs – then we can view that combination as a (super-)feature, and let it be a column in the matrix of Figure 1; the analysis described in the paper still apply. Note, however, that here we assume this super-feature construction is known initially, and in particular, this paper is *not* exploring ways to find these features – *i.e*., it is not describing machine learning tools for producing new super-features. We are also not considering multivariate approaches, where the relevance of one feature is implicitly conditioned on other features simultaneously – *e.g*., multiple regression models.

## B Exploring Other Settings

For consistency, all of the experiments in the main text used the same BD_*t*,0.05,*BH*_ biomarker discovery algorithm. However, there are many other approaches that have been used in other association studies. Here, we continue to consider only the t-test as the main statistical significance test. First, Appendix B.1 explores the obvious bootstrap method, and demonstrates that its estimate is consistently worse than our over-bound method. The other sub-appendices consider only the algorithms explicitly described in the main text. Appendix B.2 explores different options for the p-value threshold and the p-value adjustment method, to see how changing these affect the reproducibility results. This paper introduced two different approximations for the *Reproducibility Score* – uRS and oRS. Appendix B.3 explores how these approximations change as we adjust the size of the dataset *n*, and the number of iterations of running the algorithms, *k*. Finally, Appendix B.4 motivates then presents the PO score, an alternative to the Jaccard score.

### B.1 Bootstrap Sampling Method

We initially explored the standard bootstrap sampling method bRS as an overbound measure. Given a dataset *D*, this bRS method produces new datasets by drawing *n* = |*D* |subjects from *D*, with replacement: In particular, for *i* = 1..*k*, bRS(*D*, BD(·), *k*) draws a pairs of such bootstrap datasets *D*_*i*,1_ and *D*_*i*,2_ from *D*, then computes the Jaccard score of their respective BD(·)-biomarkers, and returns the average. Figures 7 and 8 show these results for continuous and SNP datasets, respectively. These experiments show, for all 25 datasets, and for both *D*^(*n*)^ and *D*^(*n/*2)^ datasets (when using all *n*, and also when using 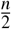 subjects), that this method is a more extreme over-bound:

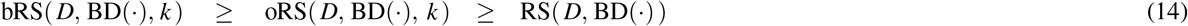

**Figure B.1.**
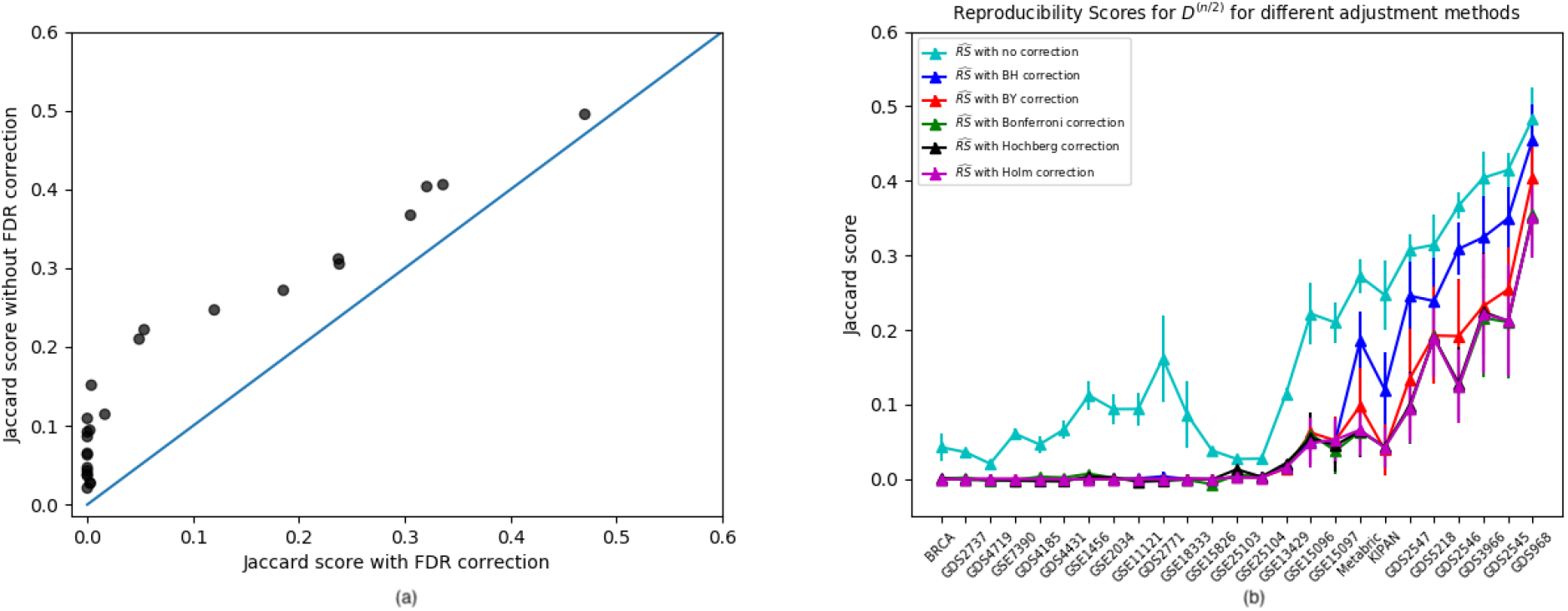
(a) Scatter plot of Reproducibility Scores for all 25 datasets: each (*x, y*) point represents the average *D*^(*n/*2)^ Jaccard scores for a single dataset (using disjoint subset pairs), where the *x*-value represents the score with MCC correction (BD_*t*,0.05,*BH*_) and the *y*-value which represents the score without MCC correction (BD_*t*,0.05,−_). Each point above the diagonal line means the MCC correction led to inferior performance for the associated dataset. (b) Reproducibility scores 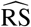 for all 25 datasets when using *D*^(*n/*2)^, for different p-value adjustment methods – *i.e*., BD_*t*,0.05,*χ*_ for 5 different FDR adjustment methods *χ*, including BH and “no”.

which means it is a less-useful measure.

### B.2 p-value adjustment methods and p-value threshold

Figure B.1(a) shows the effect of Benjamini and Hochberg (BH) MCC correction on the reproducibility scores, by comparing the average Jaccard score for the biomarkers found for a pair of complementary *D*^(*n/*2)^ datasets when using the standard MCC correction (BD_*t*,0.05,*BH*_), versus without MCC correction (BD_*t*,0.05,−_) across all datasets. We see that this MCC correction is *detrimental*, as it reduces the reproducibility scores across all datasets: While it is designed to reduce false discoveries (and hence increase precision), this may mean it is reducing recall, which collectively leads to a smaller Jaccard score.

There are many other methods for reducing MCC, in addition to Benjamini+Hochberg (BH)^49^, including: Benjamini and Yekutieli (BY)^74^, Bonferroni^75^, Hochberg^76^ and Holm^77^. Figure B.1(b) shows the results of these 5 MCC methods, as well as the “no-FDR” approach, over all 25 datasets – here showing RS with respect to the half-datasets *D*^(*n/*2)^; see Equation 8. We see again that “no-MCC” remains the best approach, and that BH is the 2nd best, followed by the others.

We also anticipate the RS score will depend on the p-value threshold used to determine the significance of each feature – *i.e*., BD_*t,τ,BH*_, for various *τ*∈ (0, 0.1). While most studies use a threshold of *τ* = 0.05, this number is fairly arbitrary. Here we explored how the RS changed with different values of *τ*. Figure B.2 shows that the reproducibility score for *D*^(*n/*2)^ appears monotonic with *τ* – within this *τ* ∈ (0, 0.1) range, larger *τ* produces higher RS.

### B.3 Changing *k*, the number of data-subset pairs drawn

Each of our approximation algorithms uses *k*, the number of data-subset pairs drawn. As we often work with large datasets, these algorithms can be very time consuming (even though they have been optimized), motivating us to explore how these algorithms scale, based on this parameter.

We therefore ran these algorithms for our largest dataset, Metabric, but varied this *k*. Figure B.3 shows that we obtained very similar results, whether we used *k* = 10, up to *k* = 50, for both uRS and oRS, for both Metabric and Metabric^(*n/*2)^. We also computed the mRS values for all 25 datasets *D* when running the algorithm for *k* = 10 versus *k* = 50, and found the difference between the two, mRS(*D*^(*n/*2)^), BD(·), k=50) − mRS(*D*^(*n/*2)^, BD(·), k=10), is very close to 0 for most cases; see Figure B.4.

### B.4 PO Score

We have used Jaccard score as our similarity measure throughout all of our experiments. This is a symmetric measure, meaning it provides information about a pair of datasets {*A, B*} where *J*(*A, B*) = *J*(*B, A*), which can be very useful when comparing the results from different experiments or evaluating the outcome when trying to replicate results from a previous study.

However, there are other options for the similarity measure that are not symmetric and can be provided together with the set of biomarkers for each dataset. One of these options is the PO score, which is used by the Zou *et al*. ^8^ meta-study: for each (ordered) pair of biomarker sets [*B*_*i*_, *B*_*j*_],

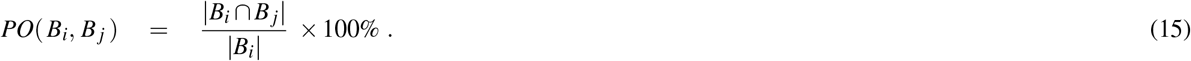

**Figure B.2.**
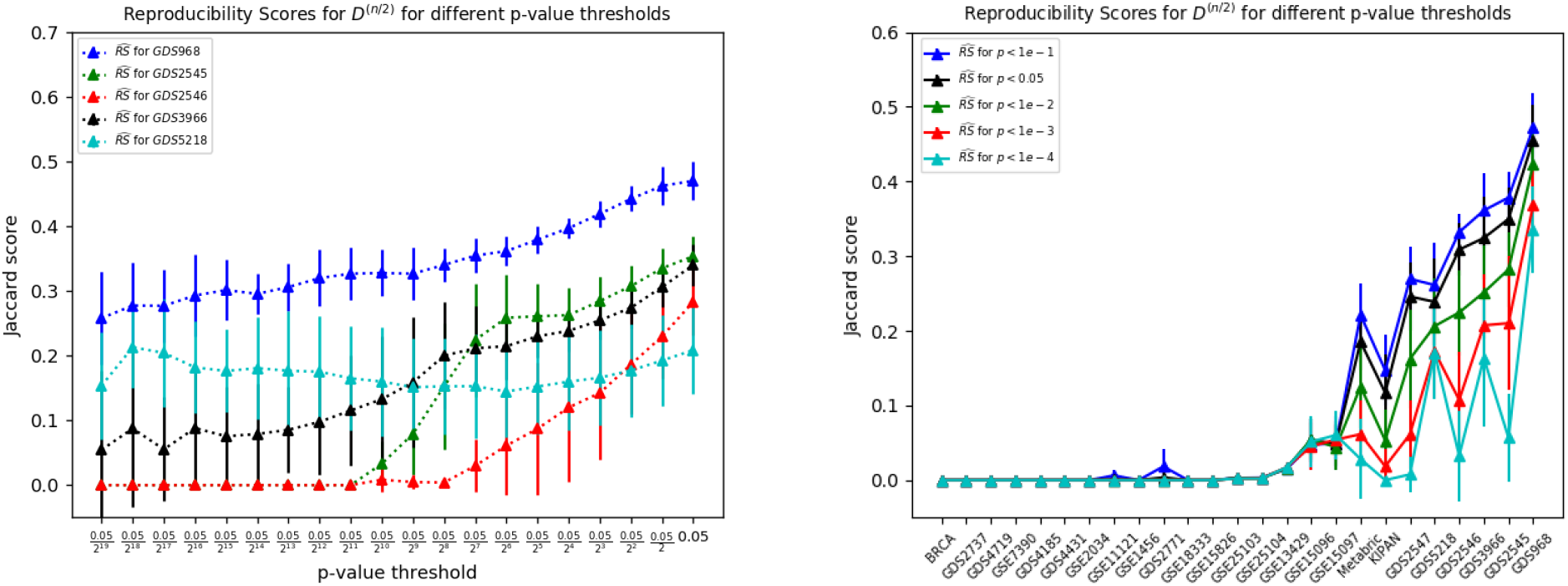
Reproducibility scores 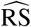 for various datasets when using *D*^(*n/*2)^ for different p-value thresholds – BD_*t,τ,BH*_, for various *τ* ∈ (0, 0.1). (left) considers 5 datasets, for a range of 20 different values of *τ*; (right) plots the Jaccard scores for all 25 datasets, for 5 different values of *τ*.

**Figure B.3.**
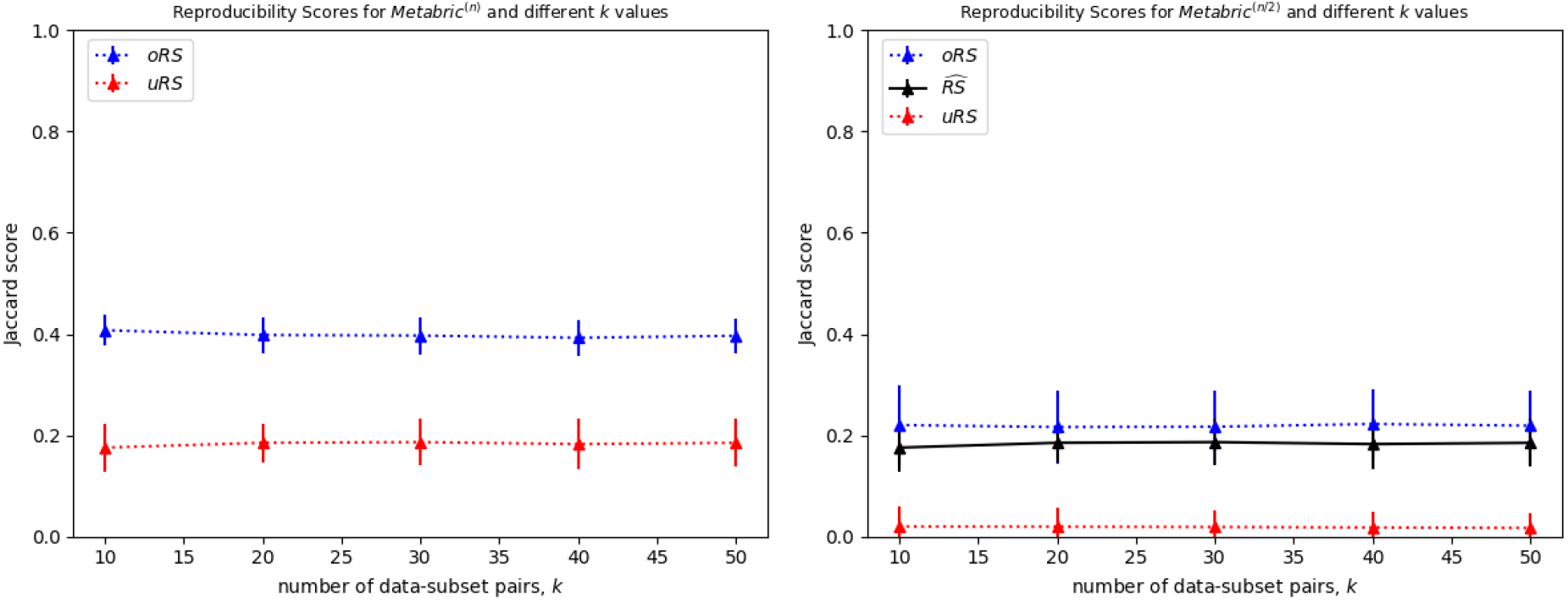
Reproducibility scores for different numbers of iterations, for the Metabric^(*n/*2)^ datasets.

Notice this is an asymmetric variant of the Jaccard score (Equation 3). (As that paper also reported the number of biomarkers found for each dataset, we could therefore recover the associated Jaccard score.) This measure can be reported with a set of biomarkers, to estimate the number of these biomarkers that should recur, if performaing another similar study.

^*RG*:^ [What else to say here?]

## C Arguments Supporting the Heuristics

The overbound and underbound algorithms, uRS and oRS, are based on some intuitions, appearing as Heuristics 7 and 9. Our empirical results, Table 1 and 2, support these claims over 25 real-world datasets – as uRS is consistently below 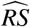, which is below oRS (as visualized by the relative heights of uRS vs 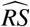 vs oRS in Figures 4(a,b)).

This appendix further motivates these heuristics. Subappendices C.1 and C.2 first provide arguments that motivate these heuristics (both focus on simple use of t-test BD_*t,α*,−_), then Appendix C.3 provides further empirical evidence that these claims hold in practice. Appendix C.4 provides additional theoretical justifications for the overlaps claims underlying Heuristic 7.

**Figure B.4.**
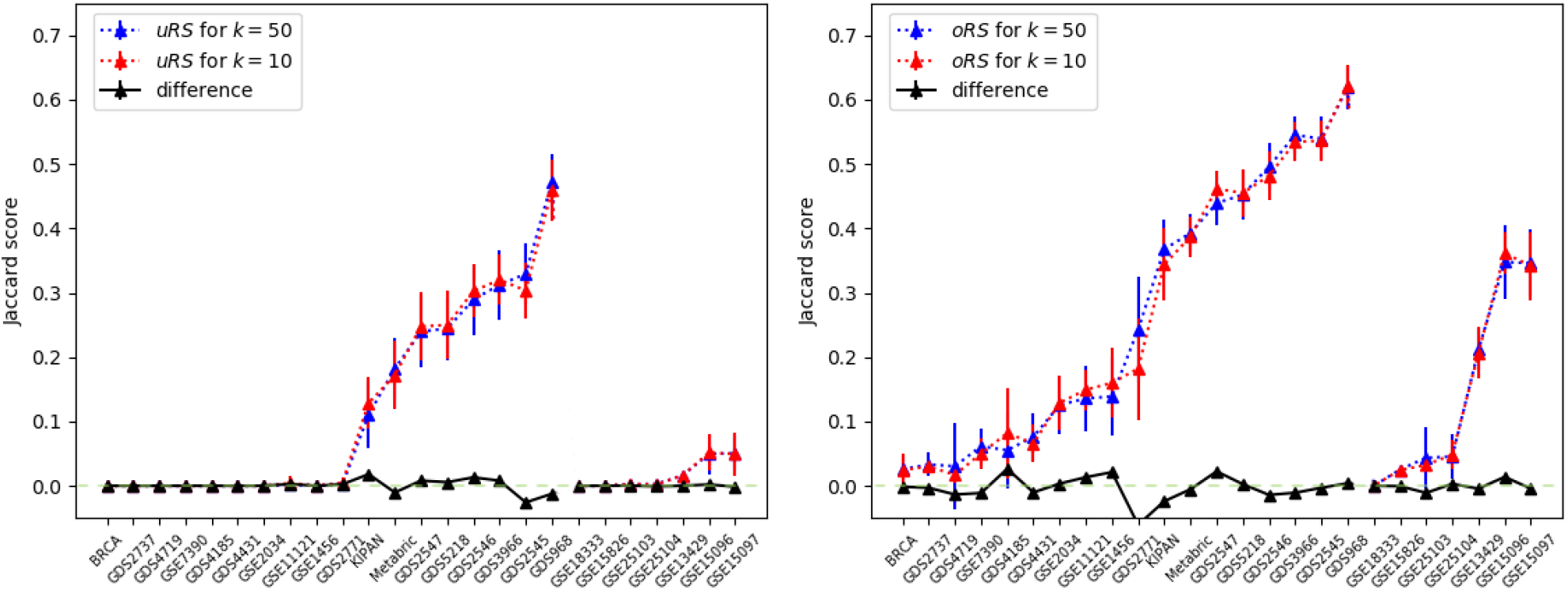
uRS and oRS values for all 25 datasets when running *k* = 10 versus *k* = 50 iterations. Note the black line, hovering around 0, is the difference between the uRS (left) values when using *k* = 10 iterations and when using *k* = 50 iterations; and similarly for the oRS values (right).

### C.1 Motivation for Heuristic 7

Here, we motivate Heuristic 7 by explaining why we expect the Jaccard score of the biomarkers found from two related datasets, to increase as the subjects forming the dataset have larger overlap. We initially consider a single feature, call it *g*, and explore a relevant statistic from these two datasets. Here, we assume that every subject of *g* associated with a positively (resp., negatively) labeled subject is drawn from a distribution with mean *µ*_+_ and variance 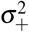 (resp., *µ*_−_ and 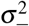). (Of course, for non-biomarkers, *µ* + = *µ* 0 and 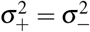 .) Each dataset has *n*^+^ positive subjects and *n*^−^ negative subjects, and these are drawn i.i.d., except that each dataset includes *r*^+^ common positive subjects and *r*^−^ common negative subjects.

For notation, let 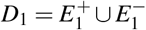 and 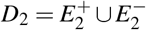 be the two datasets where 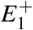 and 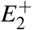 (resp., 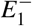 and 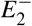) are the values of feature *g* associated with the positively (resp., negatively) labeled subjects:

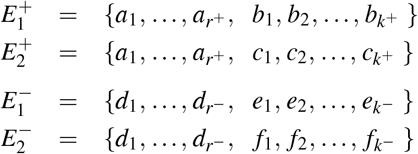

where each *a*_*i*_, *b*_*i*_, *c*_*i*_ is drawn, i.i.d., from a distribution with variance 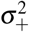, and each *d*_*i*_, *e*_*i*_, *f*_*i*_ is drawn, i.i.d., from a distribution with variance 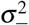.

As desired, 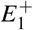 and 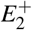 share *r*^+^ = *n*^+^ − *k*^+^ elements in common, and 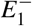 and 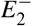 share *r*^−^ = *n*^−^ − *k*^−^ elements in common.

Now define the means:

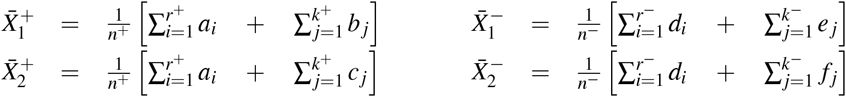

then define the joint total variance values as

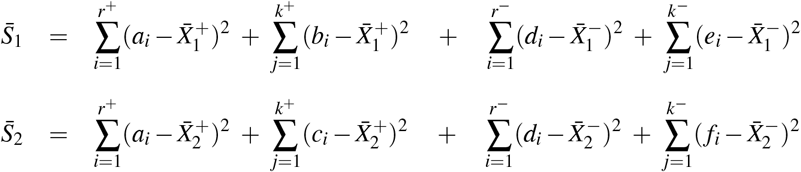

Note that 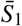 and 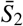 should be extremely similar, as each is the sum of *n*^+^ terms, each with expected value of 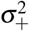, and of *n*^−^ terms each with expected value 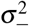. We will assume they are effectively the same value; call it 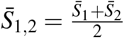

Now observe the t-statistics of each dataset:

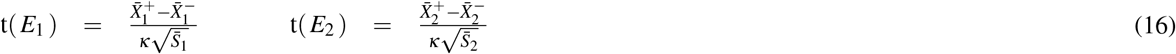

where 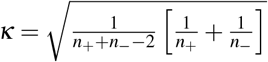, and define

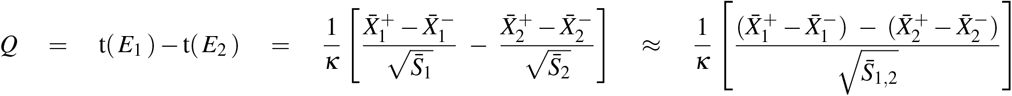

as the different between the t-statistics of these datasets. Note that the expected value *E*[*Q*] = 0. We want to show that its variance decreases as *r*^+^ or *r*^−^ increases.

We will view 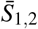 as a constant, and focus on the numerator

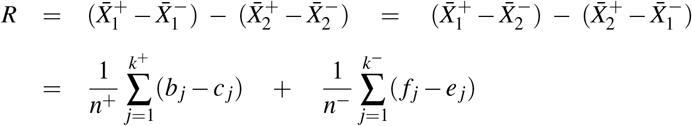

We can now observe that

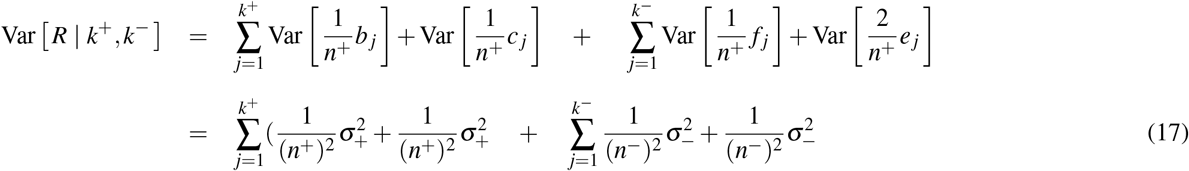

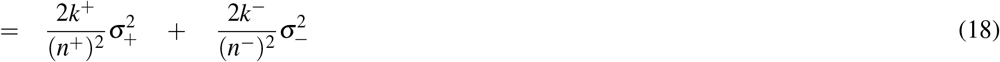

As a sanity check: if *k*^+^ = *k*^−^ = 0, then *E*_1_ and *E*_2_ are identical, meaning that they will have the same *t*-statistics, which means *Q* and hence *R* will be 0, as confirmed by this formula. Clearly this quantity **decreases** as *k*^+^ and *k*^−^ increases, which means, as *r*^+^ = *n*^+^ − *k*^+^ and *r*^−^ = *n*^−^ *k*^−^ **increase**, we expect *E*_1_ and *E*_2_ to agree more with respect to the biomarker *g* – either both agree it will be a biomarker, or both agree it will not.

Let BD(*E*_1_[*r*]), (resp., BD(*E*_1_[*r*])) be the set of biomarkers found (by our BD(·)) for *E*_1_ (resp *E*_2_), when there are *r* = [*r*^+^, *r*^−^] common elements, and let its complement *n* BD(*E*_1_[*r*]) = *F* − BD(*E*_1_[*r*]) and *n* BD(*E*_2_[*r*]) = *F* − BD(*E*_2_[*r*]) be the set of non-biomarkers, using *F* as the set of all features. The argument above suggests that | BD(*E*_1_[*r*]) ∩BD(*E*_2_[*r*]) monotonically increases with *r*, as does | *n* BD(*E*_1_[*r*]) ∩ *n* BD(*E*_2_[*r*]).

Now consider the Jaccard score, and notice

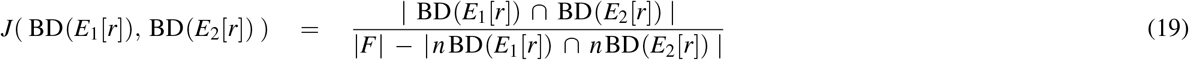

where |*χ*| is the total number of elements in the set χ. As *r* increases, we expect the numerator to increase, and the denominator to decrease, both of which means the ratio will increase.

(In general, we expect smaller values of Var [*R*] to increase the chance that *E*_1_ and *E*_2_ will agree on the status of *g*. There is one exception here: say *E*_2_ thinks that *g* is a biomarker as t(*E*_2_) is very negative, but *E*_1_ thinks it is not a biomarker, as t(*E*_1_) is near 0. Here, imagine a variant of *E*_1_ – call it 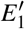 had a larger value of t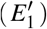; large enough that it this value meant that *g* was considered a biomarker. Here *E*_1_ and *E*_2_ disagreed about *g* with their *Q* = t(*E*_1_) − t(*E*_2_), but 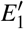 and *E*_2_ agreed about *g*, despite having a larger value of 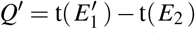. This is very unlikely, as the numerator (and hence the sign) of t(*E*_1_) is basically an empirical estimate of *µ*_1_ − *µ*_2_, which means finding a large positive value for t(*E*_1_) is unlikely unless *µ*_1_ *µ*_2_ – but this condition means it is very unlikely that t(*E*_2_) (which is another estimate of *µ*_1_ − *µ*_2_) will be very negative. That is, if *τ* if the threshold for declaring a feature to be a biomarker: *P*(t(*E*_2_) *<* −*τ* | *µ*_1_ ≫ *µ*_2_) is tiny. Similarly,

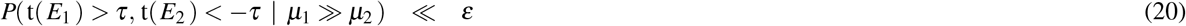

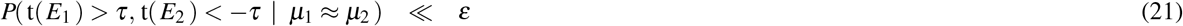

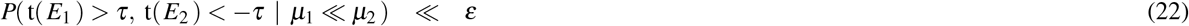

where “ ≪*ε*” means a very small probability. Equation 20 holds as t(*E*_2_) *< − τ* is unlikely in this case; Equation 21 as both t(*E*_2_) *< − τ* and t(*E*_1_) *> τ* are each unlikely in this case; and Equation 22 as t(*E*_1_) *> τ* is unlikely in this case.)

Final note: This derivation is for the general statement of Heuristic 7. Our specific oRS algorithm imposes an additional constraint – in essence, that *E*_1_ will have two copies of each *b* _*j*_ and each *e* _*j*_, and that *E*_2_ will have two copies of each *c* _*j*_ and each *f* _*j*_. The claim still holds – but with a variance for *R* that is twice as large as Equation 18: there are now only half as many *b* _*j*_s (and *c* _*j*_, *e* _*j*_, *f* _*j*_), but each is implicitly multiplied by 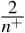 or 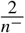, and variance goes with the square of that constant.

### C.2 Motivation for Heuristic 9

Reusing the notation from above, assume *D*_1_ and *D*_2_ are each samples of size *n*, drawn from *n*^+^ positive subjects and *n*^−^ negative subjects, independently for all of the features *F*. We also use BM to denote the set of all true biomarkers and NBM to denote the set of all true non-biomarkers.

For each gene *g*, our use of simple independent t-test means there are probabilities *p, q*(*n*) such that

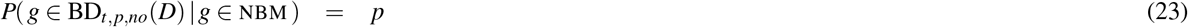

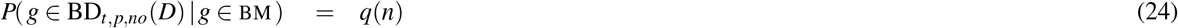

(Recall that *n* = *n*^+^ + *n*^−^ is the total number of subjects; we will insist on a constant ratio of *n*^+^ : *n*^−^ as we increase the total number of subjects.) Equation 23 is the definition of false-positive, which corresponds to the p-value of the t-test; Equation 24 is the “power” of the statistical test; we only need to know that *q*(*n*) is monotonically increasing with *n*.

Given *E*_1_[*n*] and *E*_2_[*n*] are each based on *n* draws from the 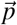 distribution, we define the expected value

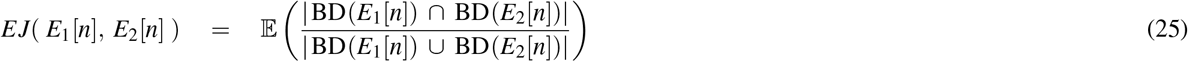

Heuristic 9 is claiming that this function is monotonically increasing as function of *n*.

To motivate this claim, we instead approximate this expectation of a ratio *EJ* (Equation 25) by the ratio of the expectations, 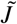 (Equation 26, below), which Lemma 1 proves is monotonically increasing in *n*.

#### Lemma 1

*Let E*_1_, *E*_2_ *each by drawn from* 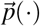 *of size n – drawing n*_+_ ∈ ℤ^+^ *instances from the joint distribution* [*p*_1,1_(·), …, *p*_*r*,1_(·)] *associated with the outcome of* +*-instances, and n*_−_ ∈ ℤ^+^ *instances from the joint distribution* [*p*_1,2_(), …, *p*_*r*,2_()] *associated with the outcome of -instances. Assuming that biomarkers are based on* BD_*t,p,no*_(*D*), *whose true positive rate is q* = *q*(*n*), *and let*

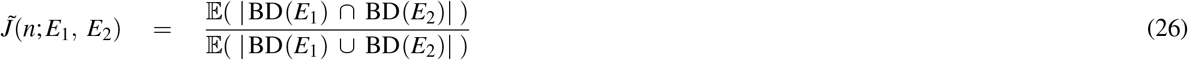

*Now assume that q* = *q*(*n*) *is increasing in n, and that q > p/*2. *Then* 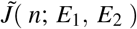 *increases as n increases*

**Proof**: We first consider a single feature *g*, and consider the likelihoods that the two sample *E*_1_ and *E*_2_ will agree: **1 What is the chance that** *E*_1_ **and** *E*_2_ **both agree that** *g* **is a biomarker?**

(1a) If *g* is a biomarker:

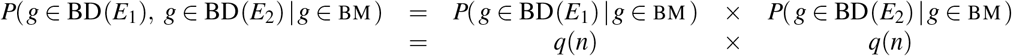

as the draws leading to *E*_1_ and *E*_2_ are independent.

(1b) If *g* is not a biomarker:

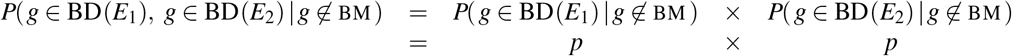

**2 What is the chance that** *E*_1_ **and** *E*_2_ **both agree that** *g* **is** *not* **a biomarker?** Here again use *n* BD(*D*) = *F* − BD(*D*) as the set of *non-biomarkers* found by BD – using *F* to represent the set of all features. Here we again deal first with the “*g* is a biomarker”, then “*g* is not a biomarker”, cases:

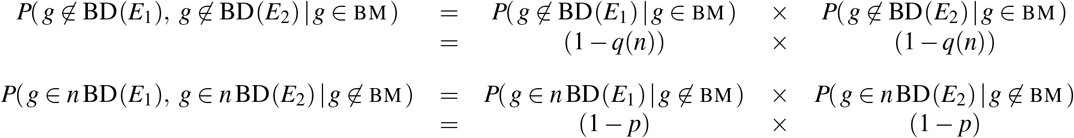

Hence, the expected size of the intersection of the biomarkers found by both samples (resp., NOT found by both) is

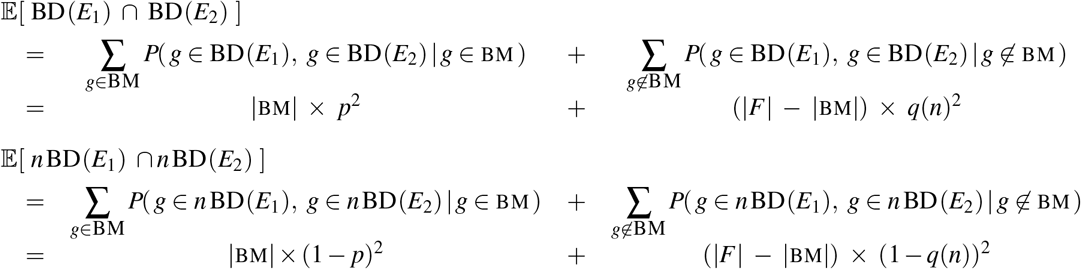

We now use the observation, from Equation 19, that BD(*E*_1_) ∪ BD(*E*_2_) = *F* − (*n* BD(*E*_1_) ∩ *n* BD(*E*_2_)), which means Equation 26 reduces to

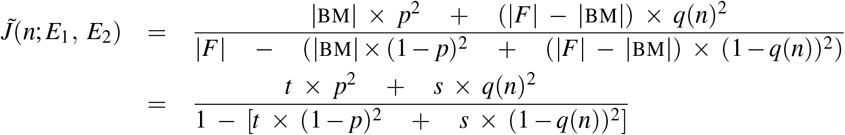

where *t* = |BM |*/*|*F* |is the fraction of biomarkers (which is typically very small), and *s* = 1 − *t* as the fraction of non-biomarkers. Also write *q* = *q*(*n*).

‘Now note that 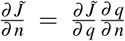, and recall that 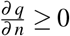. Hence, to show that 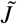 increases with *n*, it suffices to show that 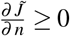.

First,

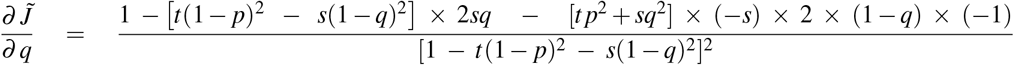

As the denominator [1− *t* (1 − *p*)^2^−*s*× (1−*q*)^2^]^2^ is positive, so we can ignore it. (Below we use ∝ to mean “same wrt sign”):

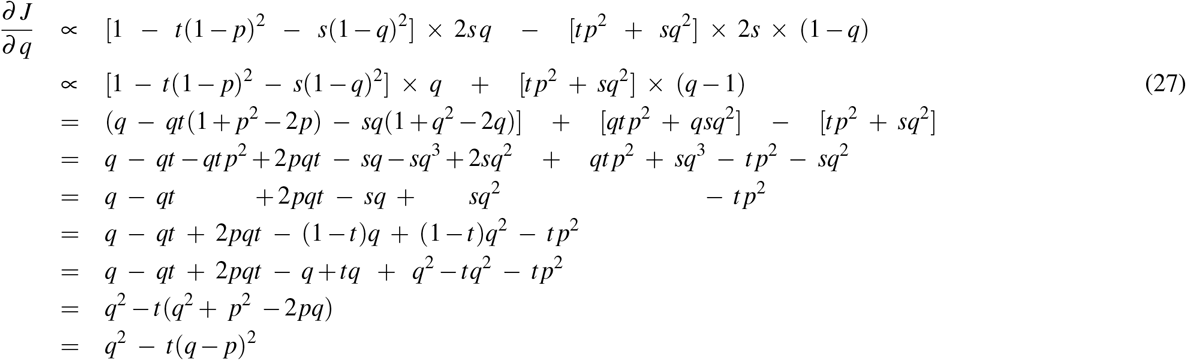

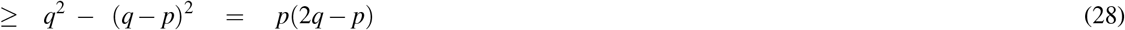

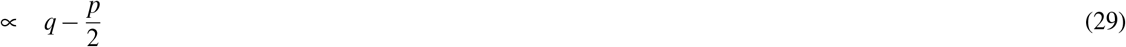

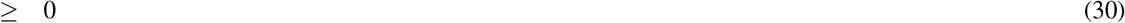

where the “∝” in Line 27 as 2*s >* 0; the “≥” in Line 28 follows from *t >* 0;, *the*”∝” in Line 29 from knowing that *p*, and hence *p/*2, is *>* 0, and the “≥” in Line 30 from our assumption that 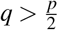. ■ (Lemma 1)

**Figure C.5.**
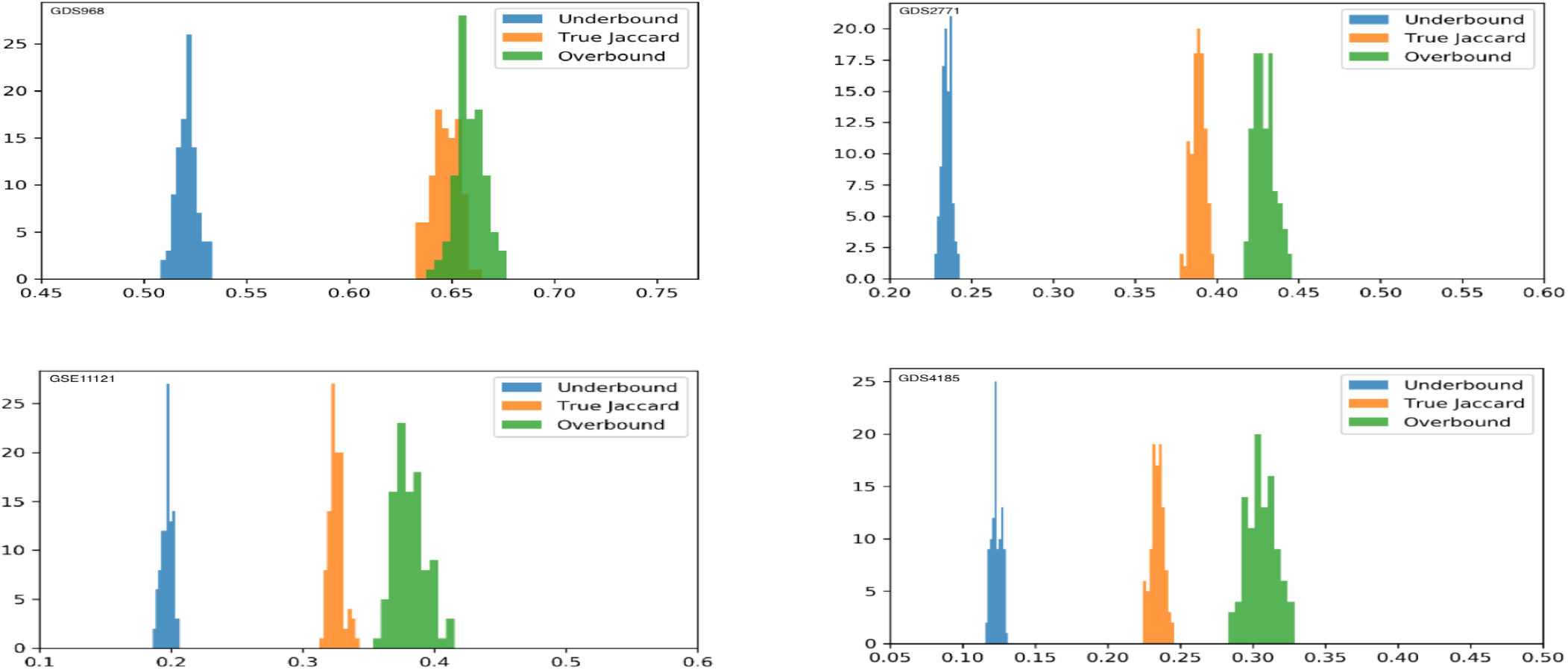
Average underbound uRS, overbound oRS and true Jaccard histograms shown for 100 pairs of simulated datasets (for the true Jaccard) and 100 simulated datasets (for underbound and overbound), with distributions for each feature/outcome based on the empirical means and variances of features of 4 datasets {GDS968, GDS2771, GSE11121, GDS4185}. Note that the x axis range is different for the different plots.

### C.3 Empirical Evidence for Heuristics 7 and 9

Our empirical results, over 25 datasets, demonstrate that algorithms based on Heuristics 7 and 9 work effectively; see Tables 1 and 2. This subappendix provides further empirical evidence by running a number of simulations, based on realistic distributions of data. In particular, we form distributions based on each of 4 real-world datasets: Γ ={GDS968, GDS2771, GSE11121, GDS4185}. (We selected these datasets to span a wide range of situations – in terms of number of subjects, number of features and the Reproducibility Scores; see Figure 5.) For each dataset, we compute the empirical mean and variance of each feature *j* and for each outcome *c*, then define *p* _*j,c*_() to be a univariate Gaussian with this mean and variance. We can then form new datasets by drawing *n*_+_(*ρ*) subjects of each *c* = + outcome and *n* _−_ (*ρ*) subjects of each *c* = − outcome, matching the size of the original *ρ* ∈ Γ dataset. (We will use *n*(*ρ*) to be the total number of elements in that dataset.)

We then ran 3 experiments on each of the 4 distributions of data. The first experiment explored the claim that “Underbound ≤ True Jaccard ≤ Overbound”. Here, we simulated 100 pairs of datasets, for each of the 4 real-world datasets, then computed the average true Jaccard score between the biomarker sets found for each pair. We also computed the average uRS and oRS score for each of these simulated datasets. Figure C.5 shows the histograms of values for each of these 4 different distributions. As expected, we see no overlap between the histograms of the underbound, true and overbound, except for GDS968. Here, we found that the true Jaccard was higher than the overbound in just 8 of the 100 cases; and the largest difference here was 0.013 (and the average difference, over these 8, was only 0.004). Figure C.6 shows the median and interquartile range for each of the 3 algorithms {uRS, true_Jaccard, oRS} for each of the 4 distributions in Γ.

The second experiment explores Heuristic 7. For each distribution (corresponding to a dataset in *ρ* ∈ Γ), for each proportion *r* ∈ {0.1, 0.25, 0.5}, we generate 100 pairs of datasets 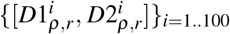 For each *i* ∈ {1..100}, we first draw *n*(*ρ*) · *r* subjects – these will be in both datasets 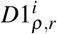 and 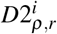. We then complete each pair by drawing the remaining *n*(*ρ*) · (1 − *r*) subjects from the underlying distribution for 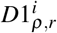, and then another *n*(*ρ*) · (1 − *r*) for 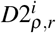. Hence, each of the *ρ*-datasets will have *n*(*ρ*) subjects, and each 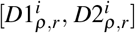 will share *r* proportion of the subjects in common.

Figure C.7 shows the Jaccard scores of each pair of datasets. We see that, as Heuristic 7 predicts, as the number of subjects in common between a pair of datasets increases, so does the average Jaccard score of the discovered-biomarker sets.

The third experiment explores Heuristic 9. In this experiment, we generate 100 pairs of datasets with *n* = |*D*| subjects (same as the associated dataset *D*), 100 pairs with *n* × 2 and 100 pairs with *n/*2 subjects, and compute the average true Jaccard score for the discovered biomarkers. Figure C.8 shows that, as Heuristic 9 predicts, as the number of subjects increases, the average Jaccard score increases as well.

Figure 4(a) earlier presented similar results, over 5 different datasets, but there taking (disjoint) subsets of the original dataset (rather than generating new subjects from the estimated distribution), and by presenting mean/variance values, rather than median/interquantile values, over 10 datbase-sizes (rather than 3). See also the black 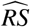 line in Figure 4(b), for yet another dataset.

### C.4 Overlap between Two Multi-Sets

This subappendix explores some relevant theoretical properties of the oRS algorithm: Recall this algorithm partitions the “doubled version” *DD* of the original dataset *D*, into 2 size-*n* sub(multi)sets, *D*1 and *D*2. Below we show that, in expectation, one-half of the elements of *D* will appear in both both *D*1 and *D*2 – *i.e*., 𝔼 [|*D*1 ∩ *D*2|] = *n/*2. We then provide a natural notion of overlap between two multi-sets, and prove that the expected overlap between 2 random draws is 0, between the *D*1 and *D*2 defined above is 1/2, and between two bootstrap samples is 1.

**Figure C.6.**
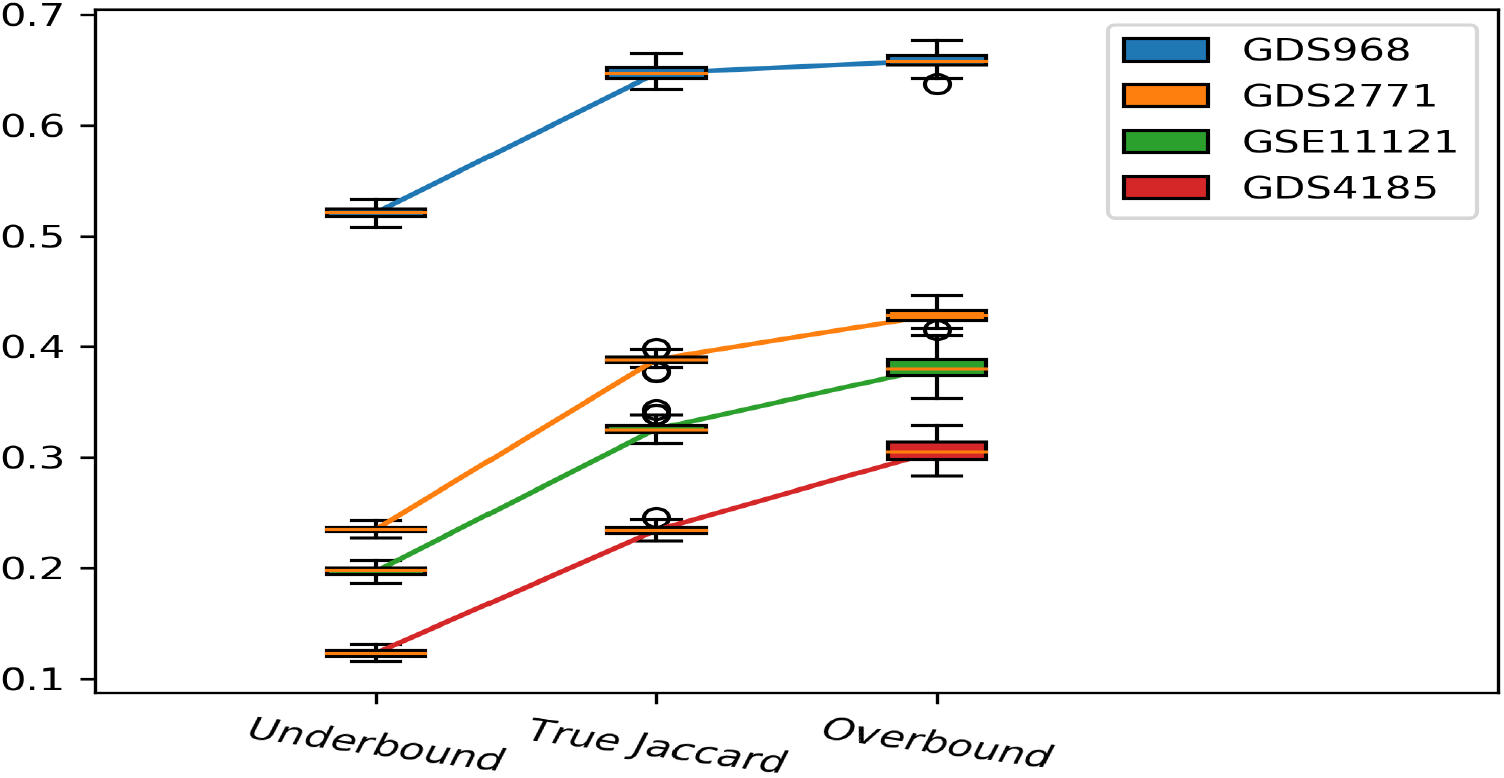
Average underbound uRS, overbound oRS and true Jaccard values for 100 pairs of simulated datasets (for the true Jaccard) and 100 simulated datasets (for underbound and overbound), with distributions based on the empirical means and variances of feature/outcome pairs of 4 datasets – using box-and-whiskers plots containing the median and IQR.

#### Expected Overlap between Multi-Sets produced by oRS

Let *DD* = {*a*_1_, *b*_1_, *a*_2_, *b*_2_, …, *a*_*n*_, *b*_*n*_} denote a set of 2*n* distinct elements. Note |*D*1 ∩ *D*2| is the number of indices *i* such that exactly one of *a*_*i*_ and *b*_*i*_ is in *D*1. We provide a formal proof below. First, to provide the intuition: If the elements drawn in *D*1 were done independently, then clearly there a 50% chance that any element of *DD* will appear in *D*1 and a 50% chance that another element will *not* appear in *D*1. If these events were independent, then the chance that *a*_*i*_ ∈ *D*1 and *b*_*i*_ ∈ *D*1 would be 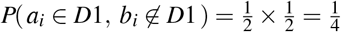. Similarly, 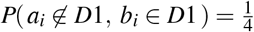. Hence, the chance that exactly one of {*a*_*i*_, *b*_*i*_} is in *A* is 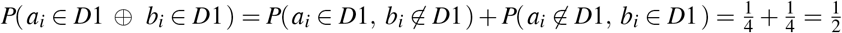.

To be precise, we need to deal with the observation that the draws from *DD* are not independent. Now let *A* be a size-*n* subset drawn uniformly from *DD* (corresponding to the *D*1 mentioned above). Let 𝒫_*n*_ denote the set of all size-*n* subsets of *DD*, equipped with the uniform distribution. For any *A* ∈ 𝒫_*n*_, let *O*_*n*_(*A*) denote the number of indices 1 ≤ *i* ≤ *n* such that exactly one of *a*_*i*_ or *b*_*i*_ is in *A*, so *O*_*n*_ : 𝒫_*n*_ → ℝ is a random variable. We view a fixed *A* ∈ 𝒫_*n*_ as the outcome of drawing a random size-*n* subset of *DD*. Note |*O*_*n*_|*/n* is the fraction of *DD* that is in both subsets.

##### Lemma 2

*Let O*_*n*_ *be defined as above. We have*

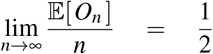

*monotonically from above*.

**Proof** Note that #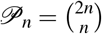, where # denotes the cardinality of a set. Suppose we are given distinct elements *Z* = {*z*_1_, …, *z*_*k*_} where *k* ≤ *n*. We compute the probability that {*z*_1_,, *z*_*k*_} ⊂ *A* where *A* ∈ 𝒫_*n*_ is a randomly chosen subset. Note that there is a bijection between size *n* subsets of *DD* containing {*z*_1_, …, *z*_*k*_} and size *n − k* subsets of *DD* \*Z* where given a subset *E* of *DD* \*Z* of size *n* − *k*, we get the corresponding subset *E* ∪ *Z* of *DD*. Thus the number of elements in 𝒫_*n*_ containing *Z* is 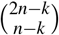, where

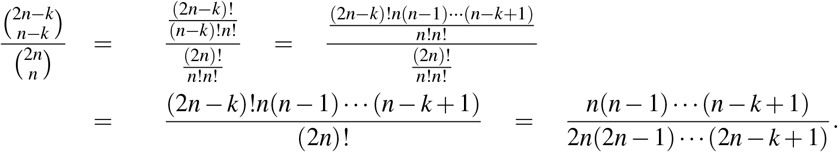

Moreover, the probability that *Z* is disjoint from a randomly chosen element of 𝒫_*n*_ is given by the same probability since for *A* ∈ 𝒫_*n*_, we have

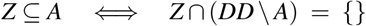

and the map *A* → (*DD* \ *A*) is a bijection from 𝒫_*n*_ to 𝒫_*n*_ since #*A* = *n*.

In particular, for a fixed index 1 ≤ *i* ≤ *n*, the probability that {*a*_*i*_, *b*_*i*_} is disjoint from a random subset of *DD* of size *n* is

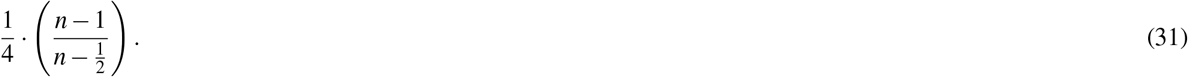

Now, let *S* _*j*_ be the indicator random variable for the event that either *a* _*j*_ or *b* _*j*_ (or both) are selected, i.e. for a set *E* ∈ 𝒫_*n*_,

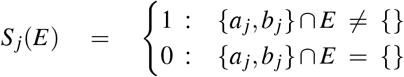

In particular, the number of distinct elements in *E*, denoted *D*_*n*_(*E*), is just given by 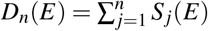. Thus by linearity of expectation and Equation 31, we have

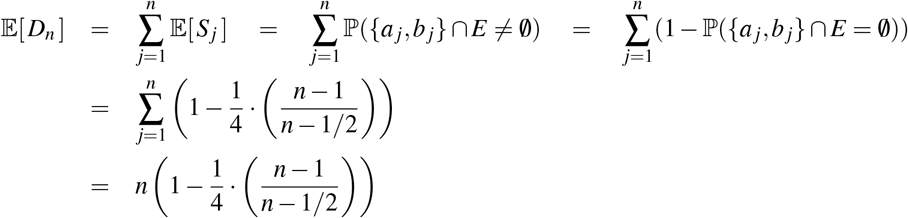

Let’s now compute the expected overlap of the sets *A* and *DD* \*A* for *A* ∈ 𝒫_*n*_. If there are *n*_1_ indices in *A* that occur exactly once, and *n*_2_ indices that occur exactly twice, then

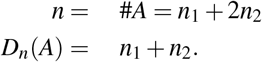

Since each index has two corresponding elements in *DD*, an index occurs in both *A* and *DD* \ *A* if and only if it occurs exactly once in *A*. In particular, if *O*_*n*_(*A*) denotes the size of the overlap of *A* and *DD* \ *A*, we have *O*_*n*_(*A*) = 2*D*_*n*_(*A*) − *n* and

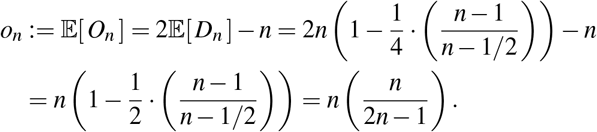

Clearly lim 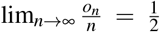, as claimed.

**Figure C.7.**
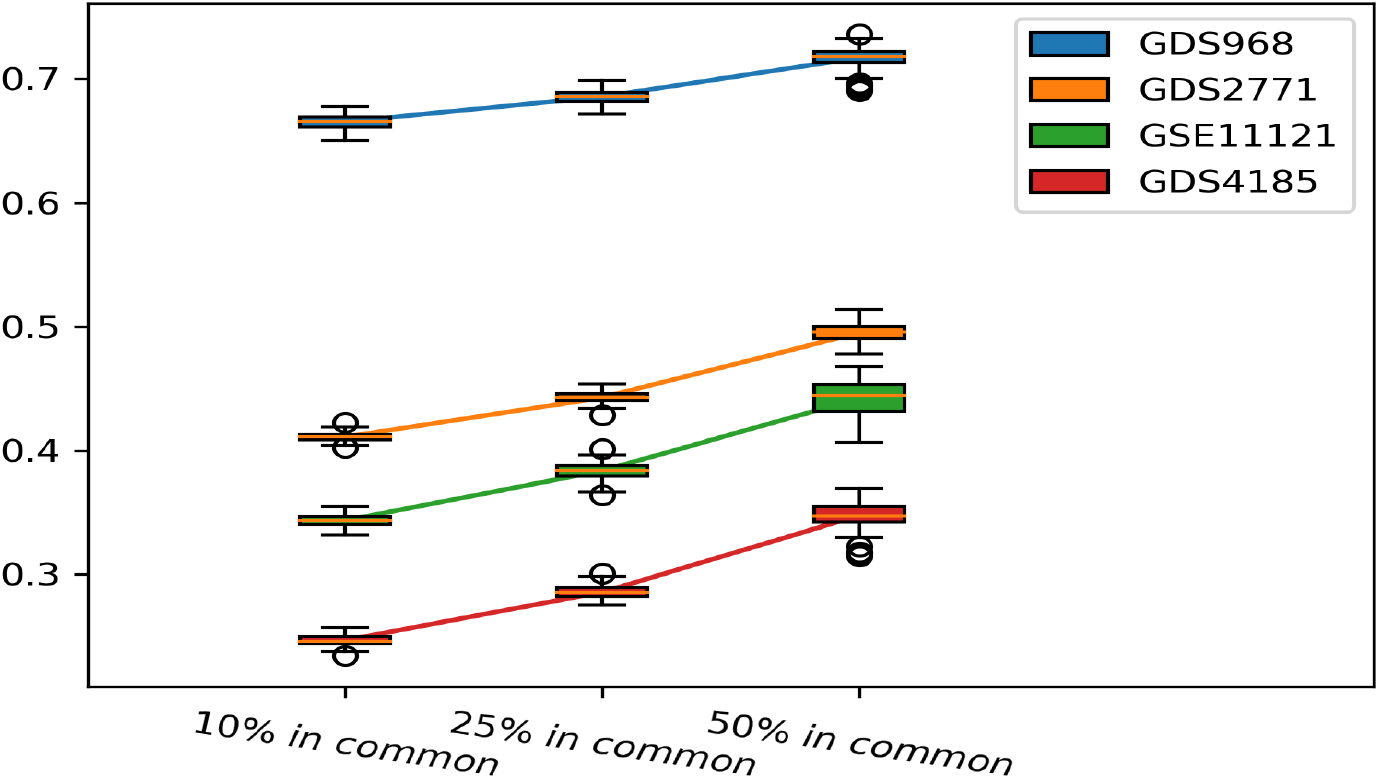
Box and whiskers plot showing the average true Jaccard values for 100 pairs of simulated datasets for 4 different distributions, in 3 settings, where each pair has 10%, 25% or 50% of the subjects in common.

**Figure C.8.**
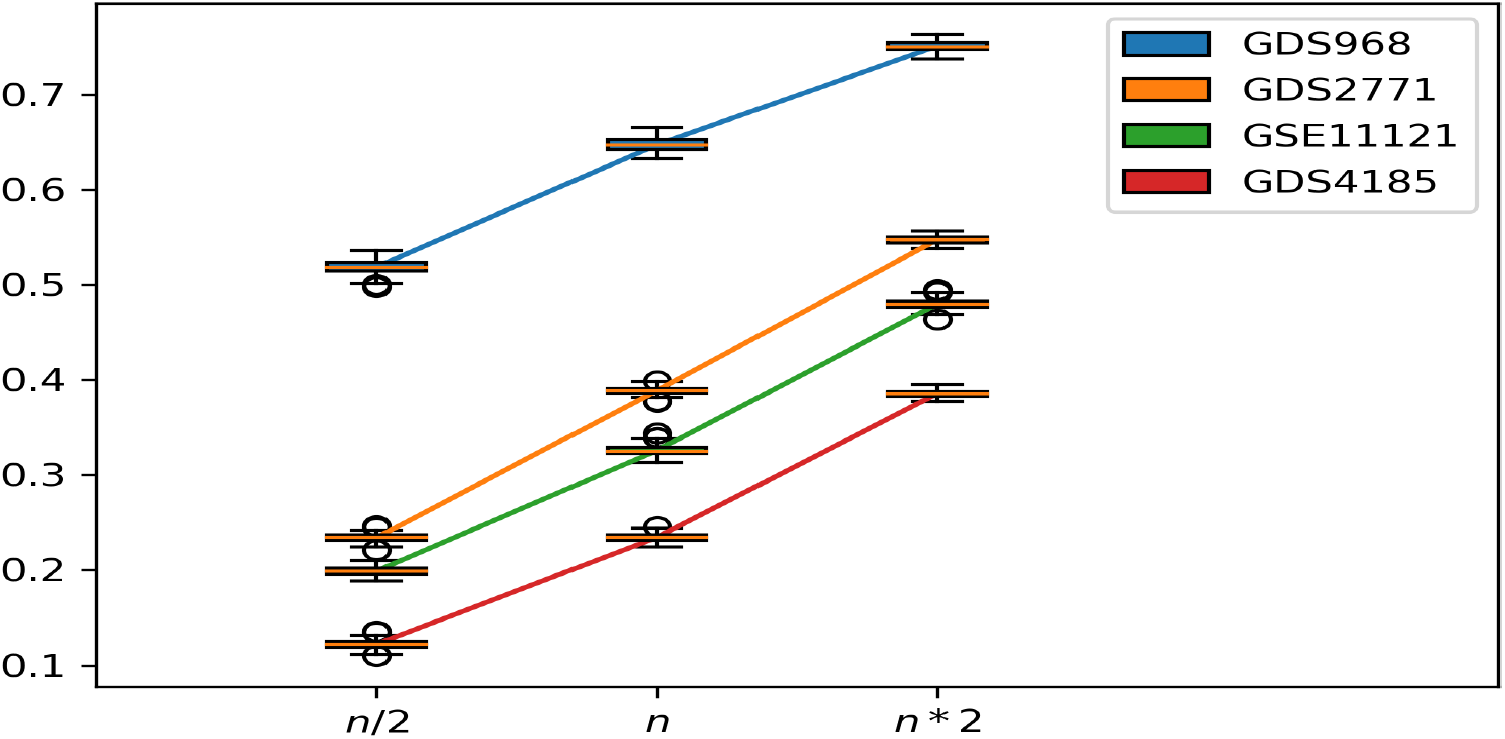
Box and whiskers plot showing the average true Jaccard values for 100 pairs of simulated datasets for 4 different distributions, and for 3 different number of subjects (x-axis) relative to the original real datasets.

#### Defining Overlap between Multi-Sets, in General

Heuristic 7 stated that the Jaccard score of biomarkers found based on two datasets, depends on the overlap between those datasets. The standard definition of overlap applies to the sets that we considered so far. To deal with bootstrap sampling, we need to extend that definition. We also show that, with this extension, we see that our oRS model has a smaller overlap that bootstrap sampling, explaining our empirical results that oRS is better than bRS.

To illustrate, let *c*(*x, S*) = the number of times that *x* appears in the (multi-)set *S*. For example, if

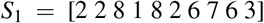

then

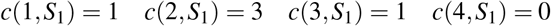

Assume |*S*_1_| = |*S*_2_|, and define:

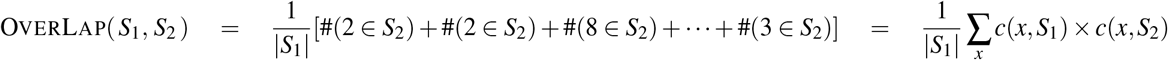

##### Lemma 3

*(a) If I*_1_ *and I*_1_ *are two size-n samples drawn independently from some continuous distribution over the reals (say from a mixture of Gaussians), then* 𝔼 [OverLap(*I*_1_, *I*_2_)] = 0

*(b) If O*_1_ *and O*_1_ *are two size-n samples produced by the oRS algorithm, from a size-n sample D, then* 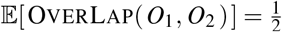.

*(c)If B*_1_ *and B*_1_ *are two size-n boot-strap samples from a size-n sample D, then* E[OverLap(*B*_1_, *B*_2_)] = 1.

**Proof**

(a) follows by realizing that the probability of any real value appearing in two size-*n* draws is effectively 0.

(b) follows from the observation that we can partition *X* = *X*_12_ ∪ *X*_1_ ∪ *X*_2_, where *X*_12_ contains elements that appear exactly 1 time in each of *O*_1_ and *O*_2_, *X*_1_ contains elements that appear 2 times in *O*_1_ and 0 times in *O*_2_, and *X*_2_ contains elements that appear 2 times in *O*_2_ and 0 times in *O*_1_. Hence,

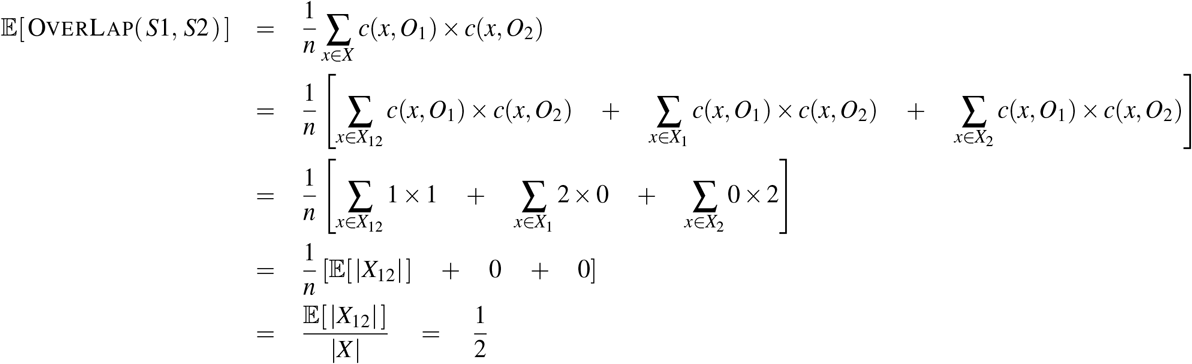

where the last line used Lemma 2, which proves that we expect *X*_12_ to contain half the elements of *X*. (c) To prove (c),

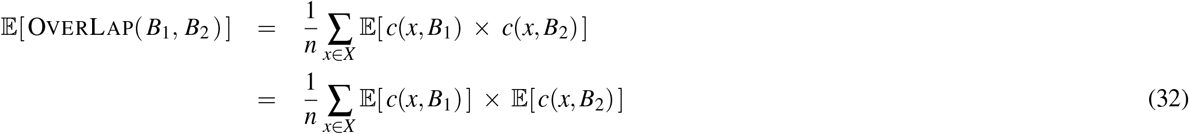

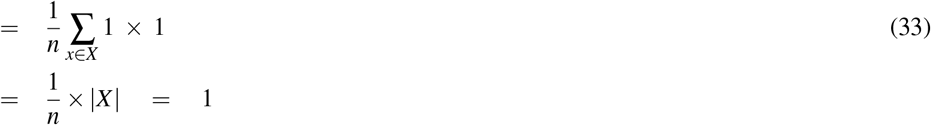

where Line 32 follows as *B*_1_ and *B*_2_ are independent bootstrap samples from *X*. To prove Line 33, note that ∑_*x*∈*X*_ *c*(*x, B*) = *n* as some value appears in each position of *B* ⇒ 𝔼 [∑_*x*∈*X*_ *c*(*x, B*)] = *n*.

Also 𝔼 [*c*(*x*_*a*_, *B*)] = 𝔼 [*c*(*x*_*b*_, *B*)] ∀*x*_*a*_, *x*_*b*_ ∈ *X* by symmetry

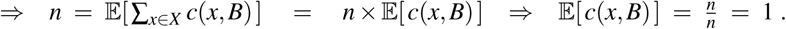

